# Beyond the greenhouse: coupling environmental and salt stress response reveals unexpected global transcriptional regulatory networks in *Salicornia bigelovii*

**DOI:** 10.1101/2020.03.17.995720

**Authors:** Houda Chelaifa, Manikandan Vinu, Massar Dieng, Youssef Idaghdour, Ayesha Hasan, Hector H. Hernandez

## Abstract

Soil salinity is an increasing threat to global food production systems. As such, there is a need for salt tolerant plant model systems in order to understand salt stress regulation and response. *Salicornia bigelovii*, a succulent obligatory halophyte, is one of the most salt tolerant plant species in the world. It possesses distinctive characteristics that make it a candidate plant model for studying salt stress regulation and tolerance, showing promise as an economical non-crop species that can be used for saline land remediation and for large-scale biofuel production. However, available *S. bigelovii* genomic and transcriptomic data are insufficient to reveal its molecular mechanism of salt tolerance. We performed transcriptome analysis of *S. bigelovii* flowers, roots, seeds and shoots tissues cultivated under desert conditions and irrigated with saline aquaculture effluent. We identified a unique set of tissue specific transcripts present in this non-model crop. A total of 66,943 transcripts (72.63%) were successfully annotated through the GO database with 18,321 transcripts (27.38%) having no matches to known transcripts. Excluding non-plant transcripts, differential expression analysis of 49,914 annotated transcripts revealed differentially expressed transcripts (DETs) between the four tissues and identified shoots and flowers as the most transcriptionally similar tissues relative to roots and seeds. The DETs between above and below ground tissues, with the exclusion of seeds, were primarily involved in osmotic regulation and ion transportation. We identified DETs between shoots and roots implicated in salt tolerance including *Sb*SOS1, *Sb*NHX, *Sb*HKT6 upregulated in shoots relative to roots, while aquaporins (AQPs) were up regulated in roots. We also noted that DETs implicated in osmolyte regulation exhibit a different profile among shoots and roots. Our study provides the first report of a highly upregulated HKT6 from *S. bigelovii* shoot tissue. Furthermore, we identified two BADH transcripts with divergent sequence and tissue specific expression pattern. Overall, expression of the ion transport transcripts suggests Na^+^ accumulation in *S. bigelovii* shoots. Our data led to novel insights into transcriptional regulation across the four tissues and identified a core set of salt stress-related transcripts in *S. bigelovii*.

## Introduction

The global population is predicted to reach 9.7 billion people by 2050 (UN, 2019). Competition between the needs for nutritional foods and sustainable biofuels production is increasing food insecurity (Kline et al., 2017; Ghosh et al., 2019). In order to meet global food and energy demands, agriculture yields need to increase from 60% to 110%, but agricultural food production is estimated to experience more than 25% drop in yield (Jenks et al., 2007; Ray et al., 2013; Qadir et al., 2014). Currently, loss of crop productivity due to increase in soil salinity affects approximately 45 million hectares of agricultural land (Boyer, 1982; Bray et al., 2000; Mittler, 2006; Roy et al., 2014).

Salinity stress is harmful to plants, affecting plant growth and yields. High concentrations of salt is toxic to plants due to osmotic stress. It affects germination processes such as seed imbibition (Khan et al., 2000; Ahmad et al., 2017), increases formation of reactive oxygen species (ROS) that cause oxidative damage to proteins and DNA (Gill and Tuteja, 2010), and inhibits the activity of nucleic acid metabolism enzymes (Gomes-Filho et al., 2008). Salt-induced stress response in plants involves intricate regulation of a diverse set of metabolic processes that play vital roles in mitigating abiotic stresses (Flowers and Colmer, 2008; Munns and Tester, 2008; Rozema and Schat, 2013).

Increased salt stress affects plant’s ability to regulate water and nutrients uptake from its environment while avoiding excessive accumulation of salt ions in its tissue. Osmotic stress induces expression of genes encoding antioxidants, ion channels (Wu et al., 2009), potassium transporters, vacuolar or plasma membrane Na^+^/H^+^ (Zhang et al., 2008; Oh et al., 2009; Fan et al., 2013), vacuolar pyrophosphatases (Silva and Gerós, 2009), and proteins involved in defense functions and signal transduction (Zhu, 2002). In addition, plants synthesize a set of organic solutes, such as sugars, amino acids, and glycine betaine (GB), and accumulate inorganic ions in vacuoles to help maintain turgor (Hasegawa et al., 2000; Khan et al., 2000; Flowers et al., 2010). The synchronous process of gene expression and metabolite production allows plants to adjust to varying salinity stresses and thus maintains a positive turgor pressure.

To date, the molecular mechanisms of salinity response in crop plants are poorly understood due to the fact that there is a paucity of halophyte plant genomes and global transcriptome studies focusing on response to salt stress (Flowers et al., 2015). Understanding the mechanisms underlying salt tolerance and investigating the use of halophytes as food sources are key to addressing the predicted shortfalls in food production.

Developing halophyte crops for use in semi-arid regions where soils suffer from high salinity and water scarcity is a vital part of the effort to increase the mix of food crops. The *Salicornia* genus, commonly known as glasswort or sea asparagus, is a widespread extremely salt-tolerant succulent annual herb from the Chenopodiaceae family (Kadereit et al., 2006; Kadereit et al., 2007). It grows along seashores, colonizing new mud flat areas through prolific seed production (Glenn et al., 1997; Glenn et al., 1999). It is a leafless, succulent small-seeded saltmarsh plant considered a promising saline water crop (Glenn et al., 1992; Glenn et al., 1997; Zerai et al., 2010).

*S. bigelovii* shows great promise as a semi-arid saline crop for conserving freshwater, providing food, fodder, and producing biofuels (El-Mallah et al., 1994; Anwar et al., 2002; Warshay et al., 2017). It displays high seed yield, which can contain greater than 30% edible oil, and high biomass production under seawater irrigation (Glenn et al., 1998). Currently, *S. bigelovii* is commercially cultivated as a minor vegetable for the United States and European fresh produce markets. It is one of the world’s first terrestrial crops produced exclusively under seawater irrigation in several large projects (Glenn et al., 1992; Glenn et al., 2013).

Generally, transcriptomic studies of abiotic stress response in plants are performed in the laboratory or in greenhouses under strictly controlled conditions. These studies center on understanding plant response under a singular condition using a single snapshot in time to investigate the differential transcriptome in response to saline stress. In addition, they concentrate on specific plant tissues, such as root and shoots (Fan et al., 2013). Unfortunately, this approach does not reflect the real-life conditions that plants face in the field (Atkinson and Urwin, 2012; Ramegowda and Senthil-Kumar, 2015; Pandey et al., 2017). Conducting transcriptome studies under controlled conditions can overlook the combined effects of other environmental factors, such as heat stress and salinity, in modulating the global transcriptome response of plants to abiotic stresses (Pandey et al., 2017).

There are several key adaptive traits specific to *S. bigelovii* that allow it to germinate, mature, and complete its life cycle under saline conditions (Yuan et al., 2019). The effects of salt stress are felt in all growth phases of *S. bigelovii* throughout multiple metabolic, molecular, and physiological processes (Rivero et al., 2014; Negrão et al., 2016; Salazar, 2017).

Here, we (1) generated global tissue-specific (flowers, roots, seeds and shoots) *S. bigelovii* transcriptomes from plants grown in a test facility under varying environmental conditions and saline aquaculture effluent irrigation, (2) identified transcripts differentially expressed in *S. bigelovii* tissues, (3) characterized a core set of salt stress-related transcripts, and (4) studied *S. bigelovii*’s metabolic responses implicated in salt stress adaptation and osmolyte production induced by growth under saline aquaculture effluent. The understanding of global transcriptome response to salt stress is key if we are to elucidate the underlying mechanisms of salt stress response in both halophytes and other crop plants. These data represent the first global transcriptome of salinity stress response in *S. bigelovii*. One novel finding in our study is the identification of previously unreported tissue expression patterns of salt stress response transcripts, such as HKT6m SOS1, and NHX. These results offer novel insights into the modes of transcriptional and metabolic regulation that can potentially be exploited to facilitate the development and selective breeding of crops with increased tolerance to salt stress.

## Materials and Methods

### Seawater Energy and Agriculture System field station

The Seawater Energy and Agriculture System (SEAS) pilot facility (Supplementary Figure S1) is an integrated aquaculture, halo-agriculture, and mangrove silviculture system located at the Khalifa University, Masdar City Campus, Abu Dhabi, United Arab Emirates. The facility uses sea water aquaculture ponds to breed fish and shrimp with the aquaculture effluent flooding *S. bigelovii* fields twice per day, mimicking tidal flows. The SEAS pilot facility experiences large annual fluctuations in environmental conditions with average maximum air temperature of 25.9°C in January to 44.1°C in August and with the majority of the rainfall occurring between December and March (Statistics Center, 2018). Sea effluent water was analyzed by Ion chromatography IC-5000 (Dionex ICS-5000, ThermoFisher) with separator column (3×100) to analyze cations and ions concentration.

### Plant material and RNA extraction

*S. bigelovii* cultivar ‘Boca Chica’ (Arizona University, Tucson, Arizona, USA) seeds were selected and grown under environmental conditions at the SEAS pilot facility. Four *S. bigelovii* plant tissue, namely flowers (F), roots (R), seeds (G), and shoots (S) (Supplementary Figure S2), were collected from randomly selected individual plants from the same field area, immediately frozen on site in liquid nitrogen, transferred to the laboratory, and stored at −80°C for downstream analysis.

Total RNA was extracted from three biological replicates of each of the four tissues using the RNeasy Plant Mini kit (Qiagen, Venio, Netherlands). The purity of RNA samples (RNA Integrity Number > 8) was confirmed using a Bioanalyser RNA Chip 2100 (Agilent Technologies, Santa Clara, CA, USA) and quantified using a Qubit 4 Fluorometer (Invitrogen, Carlsbad, CA, USA).

### cDNA library preparation and next-generation sequencing (NGS)

All cDNA libraries were constructed using the TruSeq Stranded mRNA Library Prep Kit (Illumina, San Diego, CA, USA) following the manufacturer’s protocols. Enrichment of mRNA from total RNA was performed using poly-T attached magnetic beads followed by enzymatic fragmentation and cDNA synthesis using Superscript II Reverse Transcriptase (Thermo Fisher, Waltham, MA, USA). cDNA samples were purified using AMPure XP beads (Agencourt Bioscience, Beverly, MA, USA) followed by adapter and barcode ligation, A-tailing, and amplification as recommended by the manufacturer. The resultant cDNA libraries were subjected to 250 bp paired-end sequencing using an Illumina Hi-Seq 2500 platform (Illumina, San Diego, CA, USA).

### Global analysis, *de novo* assembly, and visualization of RNA reads

Transcriptome reads data were processed to remove adapter sequences, low quality reads (QV < 30 Phred score), and reads of length below 36 bp using the program Trimmomatic v0.32 (Bolger et al., 2014). The quality of the trimmed reads was assesses using FastQC 0.11.4 (Andrews, 2010). Quality control of the raw data resulted in retaining high-quality transcriptome data of 190,227,720 paired-end reads for all tissues. We performed *de novo* assembly of the trimmed reads using the assembler Trinity (V2.2.0) (Grabherr et al., 2011). Transcript quantification was done via the utility script ‘align_and_estimate_abundance.pl’ bundled in the Trinity toolkit using default values with the alignment and estimation methods set to Bowtie2 and RSEM, respectively. Assembled transcripts with a minimum length of 200 bp and expression levels of 5 (FPKM) were retained for downstream analysis. The non-redundant transcripts were further clustered using CDHIT-EST at 95% identity.

Unsupervised statistical analysis and two-way hierarchical clustering was used to plot global gene expression patterns using the JMP Genomics software (SAS Institute, Cary, NC, USA) using log-transformed normalized transcript abundance values. All RNA sequencing data for the replicates and four tissues (R, S, F, and G) have been deposited at the NCBI in the Short Read Archive database (Bioproject ID: PRJNA607385, Biosample ID: SUB6825665).

### Differential gene expression analysis

FPKM (Fragments Per Kilobase of transcript per million mapped reads) values were calculated to measure the expression level of each assembled transcript sequence. This method eliminates the influence of varying transcripts lengths and sequencing depth on the calculation of transcript expression. Differential expression analysis was performed using count data and DESeqv1.8 based on the negative binomial distribution model. Significant differential expression was inferred based on a false discovery rate (FDR) of 5% and a log2 fold change ≥ 2.

### Functional gene annotations, gene ontology, and pathway analysis

Assembled transcripts with a minimum length of 200 bp and expression levels (FPKM) of 5 were used for downstream analysis using Blast2GO v4.1.9 (Götz et al., 2008) to identify the GO terms associated with biological processes (BPs), molecular functions (MFs), and cellular components (CCs) (https://www.blast2go.com/blast2go-pro). All putative transcripts were searched against the NCBI database (nr) (ftp://ftp.ncbi.nih.gov/blast/db/29-02-2015), UniProt database (Consortium, 2018), Kyoto Encyclopedia of Genes and Genomes (KEGG) (Kanehisa and Goto, 2000), Gene Ontology (GO) (Ashburner et al., 2000; The Gene Ontology Consortium, 2018), EggNOG 5.0 (Huerta-Cepas et al., 2016) and InterProScan (Madeira et al., 2019) using Basic Local Alignment Search Tool (BlastX) with an E-value cut-off set to 10^−3^ and a minimum number of hits of 10. Only the top hit from the BlastX search were used for downstream analysis.

### Quantitative PCR

Nine transcripts implicated in salt stress response were selected for quantitative real-time PCR (qRT-PCR) validation. The qRT-PCR was performed as previously described (Wang et al., 2016). First-strand cDNA was synthesized using a PrimeScript RT reagent kit followed by qRT-PCR using SYBR Green qPCR Master Mix on a StepOnePlus Real-Time PCR instrument (Thermo Fisher Scientific, Waltham, MA, USA) following the manufacturers’ protocols. *GADPH* was used as the reference gene in accordance with previous methods (Hao et al., 2014). The transcript primers used are listed in Supplementary Table S2. Relative gene expression levels were calculated using the 2^−ΔΔCT^ method (Livak and Schmittgen, 2001), and the data are presented as the mean□±□SD from three independent biological replicates.

## Results

### Environmental conditions at SEAS field station during tissue sampling

The SEAS field station, located in Masdar City, Abu Dhabi, is designed to assess environmental parameters that can affect crops production scale systems for saline agriculture. The meteorological parameters at the SEAS field station track the annual fluctuations typically experienced in Abu Dhabi (Supplementary Figure S3A (www.Average-Weather-at-Abu-Dhabi-International-Airport-United-Arab-Emirates)). Salinity of the aquaculture effluent used to irrigate *S. bigelovii* fields ranged from 32.4 g L^-1^ in March to 42.5 g L^-1^ in August. The constant relative low water salinity in the first period of growth (March-April 2017) as compared to the high salinity in the second period of growth (July-August) can be attributed to the rainy days in late winter and early spring as well as higher mean air temperature and evaporation in summer (Supplementary Figure S3C). The effluent sea water ion content over the measured period varied slightly with salinity increase. In general, we observe a trend of high concentration of Na^+^ (11.7 g L^-1^ to 16.9 g L^-1^), K^+^ (0.53 g L^-1^ to 0.65 g L^-1^), and Cl^−^ (18.7 g L^-1^ to 21.9 g L^-1^) (Supplementary Table S1).

A total of twelve *S. bigelovii* plants tissue samples representing roots (R1, R2, R3), shoots (S1, S2, S3), flowers (F1, F2, F3), and seeds (G1, G2, G3) tissue were collected between March 2017 and August 2017 from plants grown at the SEAS facility. Shoots and roots were collected in March 2017 from the same plant, while flowers and seeds were collected in July and August 2017, respectively. These tissues were used to investigate the transcriptome of *S. bigelovii* plants in response to growth under environmental conditions.

### Sequencing and *de novo* assembly of the *S. bigelovii* transcriptome

Differential expression analysis without a reference annotated genome is challenging. Nowadays however, this analysis is possible using data with sufficient sequencing depth and *de novo* transcriptome analysis approaches (Pucker and Schilbert, 2019). Deep *S. bigelovii* transcriptomic data was generated and subjected to *de novo* transcriptome analysis. A total of 226,274,905 raw reads for all the libraries processed were generated using the Illumina HiSeq platform. Quality control using FastQC (Andrews, 2010) resulted in retaining 190,227,730 reads (88%) with GC content of 45.28% (Table 1). All sequencing data generated have been deposited in the NCBI database under the NCBI SRA accession number PRJNA607385.

**Table 1.** Sequencing and assembly statistics of *S. bigelovii* transcriptome.

| Organs | number of trimmed reads |
| --- | --- |
| Flowers | 43,459,494 |
| Shoots | 47,133,647 |
| Roots | 48,398,048 |
| Seeds | 51,236,541 |
| **Total reads** | 190,227,730 |
| **Total assembled transcripts** | 643,752 |
| **Percent GC (%)** | 45,28 |
| **Smallest (bp)** | 200 |
| **Average (bp)** | 557 |
| **N50 (bp)** | 722 |

The roots, shoots, flowers, and seeds libraries produced 48,398,048, 47,133,647, 43,459,494, and 51,236,541 reads (quality scores greater than Q20), respectively (Figure 1). The *de novo* assembly yielded 643,752 high-quality transcripts of length between 200 bp and 2,627 bp with an N50 of 722 bp (Table 1, Figure 1A). The redundant transcripts represented less than 20% of the total reads as analyzed by CD-HIT-EST (Fu et al., 2012) using a 95% identity cutoff.

**Figure 1.**
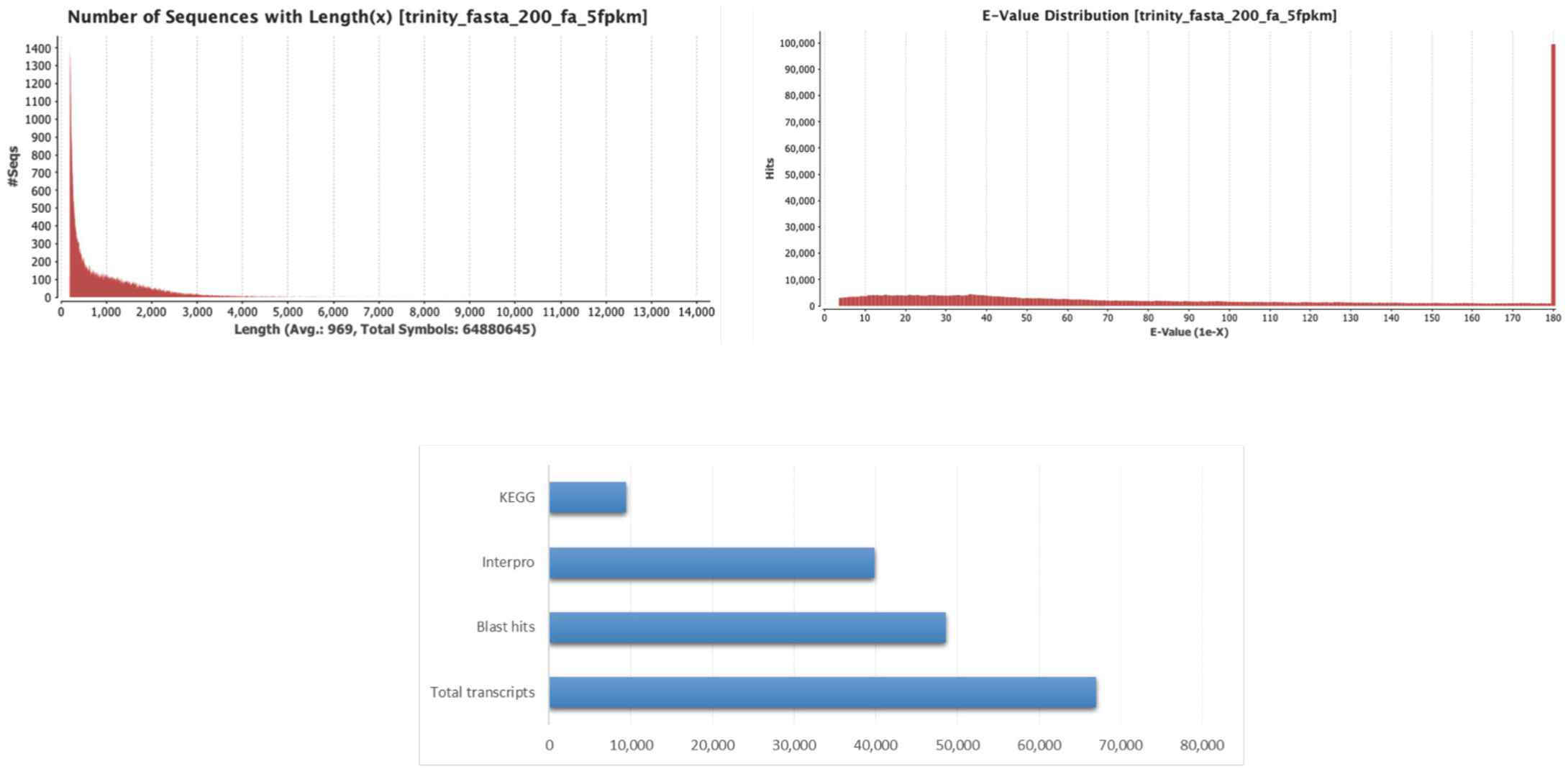
Functional annotation of *S. bigelovii* transcriptome. **(A)** Global sequence distribution transcripts length, **(B)** *E*-value distribution of BLAST hits for each unique sequence against the N database, (**C)** Distribution of Blast2GO three step processes including KEGG, InterPro, and BLAST hits of total number of assembled transcripts.

### Functional annotation and classification

Assembled transcripts with a minimum length of 200 bp and expression levels of 5 (FPKM) in at least 2 biological replicates per tissue type were retained for downstream analysis. The resultant 66,943 transcripts were submitted for annotation using the Blast2GO program. Based on this criteria, 48,622 transcripts (72.63%) were matched in the GO database, 39,882 (59.56%) in the InterPro database, and 9,424 (14.07%) in the KEGG database (Figure 1C).

Since there are no reference *Salicornia* species genomes available in public databases, we compared the *S. bigelovii* transcripts against the complete NCBI database using BLAST. The total number of BLAST top hit sequences was 66,943, with 48,613 of these sequences assigned to a known species (Figure 2). Assessment of the distribution of the top-hit species from this analysis showed that 18,936 transcripts (38.94%) had high homology with genes from *Beta vulgaris maritima*, followed by *S. oleracea* (9,692 transcripts, 19.93%) and *Vitis vinifera* (146 transcripts, 3.03%), while 17,029 transcripts had high homology with sequences from other organisms (Figure 2). A total of three fungal endophytes, namely; *Hortaea werneckii* (4,021 transcripts, 6%), *Rachicladosporium* sp. (3,096 transcripts, 4.62%), and *Acidomyces richmondensis* (592 transcripts, 0.8%) were detected.

**Figure 2.**
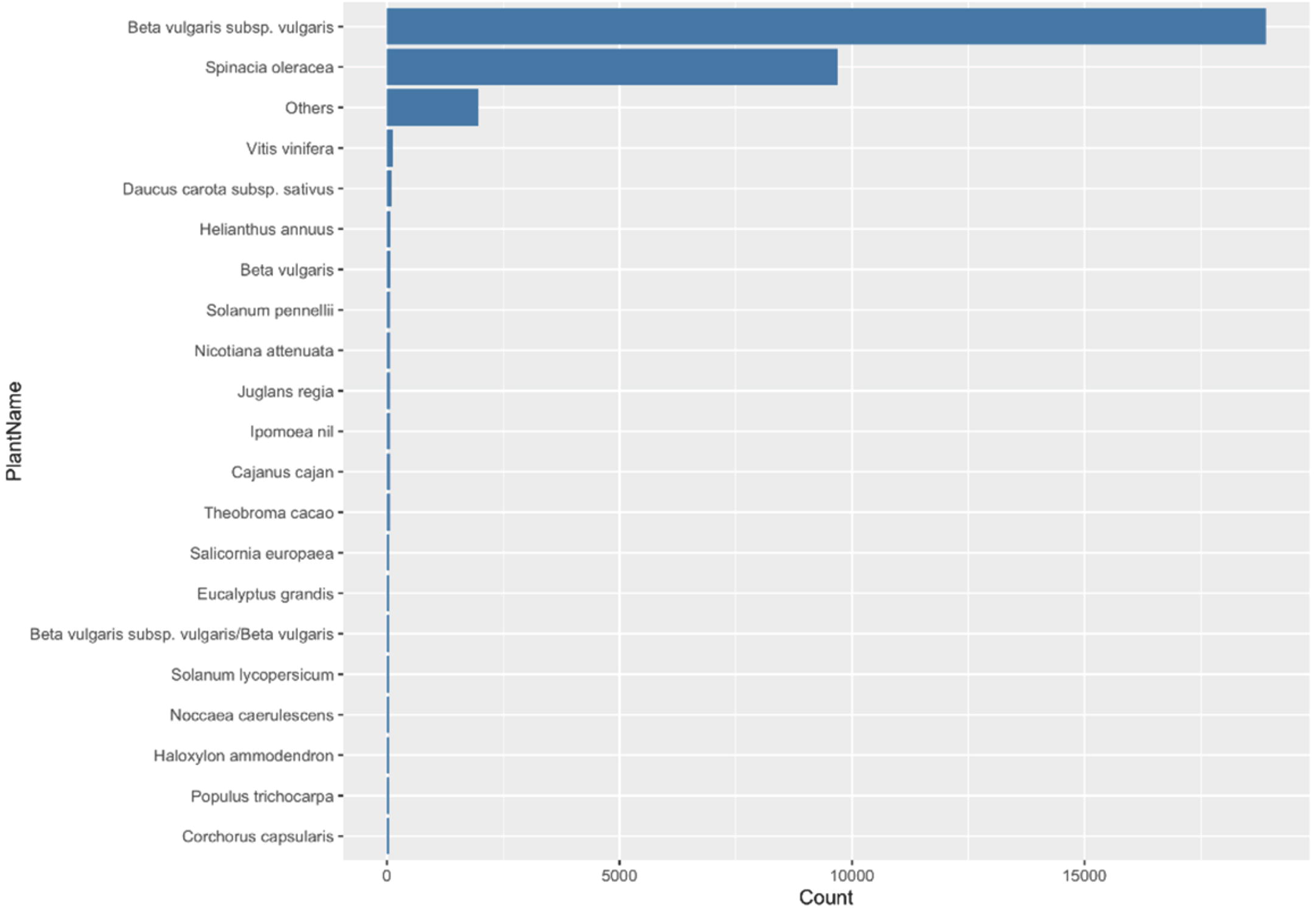
Annotation of the *S. bigelovii* transcriptome. Species distribution of the top 20 plant species based on BlastX alignments against the NCBI nr database.

All transcripts identified were subject to the GO classification. A total of 66,943 Trinity-assembled *S. bigelovii* transcripts were assigned GO terms. A total of 133,971 annotations were identified from the BLAST results (Figure 3A). Based on sequence homology with genes of known function, these transcripts could be assigned to one or more ontologies, with 35% of transcripts assigned to a biological process (BP), 26% of transcripts assigned to a cellular component (CC), and 39% of transcripts assigned to a molecular function (MF) category (Figure 3). For BPs (Figure 3B), a maximum number of transcripts (20,250 transcripts) were associated with plant metabolic processes followed by cellular processes (17,715) and single-organism process (12,560). Additional transcripts identified implicate response to stimulus (3,145), signaling (1,081), developmental process (976), reproductive process (579), immune system process (95), rhythmic process (20), locomotion (24), biological adhesion (20), and cell killing activity (6). For MFs, transcripts were associated with catalytic activity (18,089) and binding (16,613) followed by genes of transporter (2,637) and structural molecular activity (2,308). Other transcripts related to MF included molecular function regulation (425), transcription factor to nucleic acid binding (403), transcription factor to protein binding (157), electron carrier nutrient reservoir (64), and metallochaperone activity (5). CCs implicated (Figure 3B) include cell part (15,344), membrane function (12,005), and membrane part (9,965) followed by organelle part (5,673) and extracellular region (569). Other transcripts have been annotated as implicated in cell junction (197), nucleoid (157), and virion (91).

**Figure 3.**
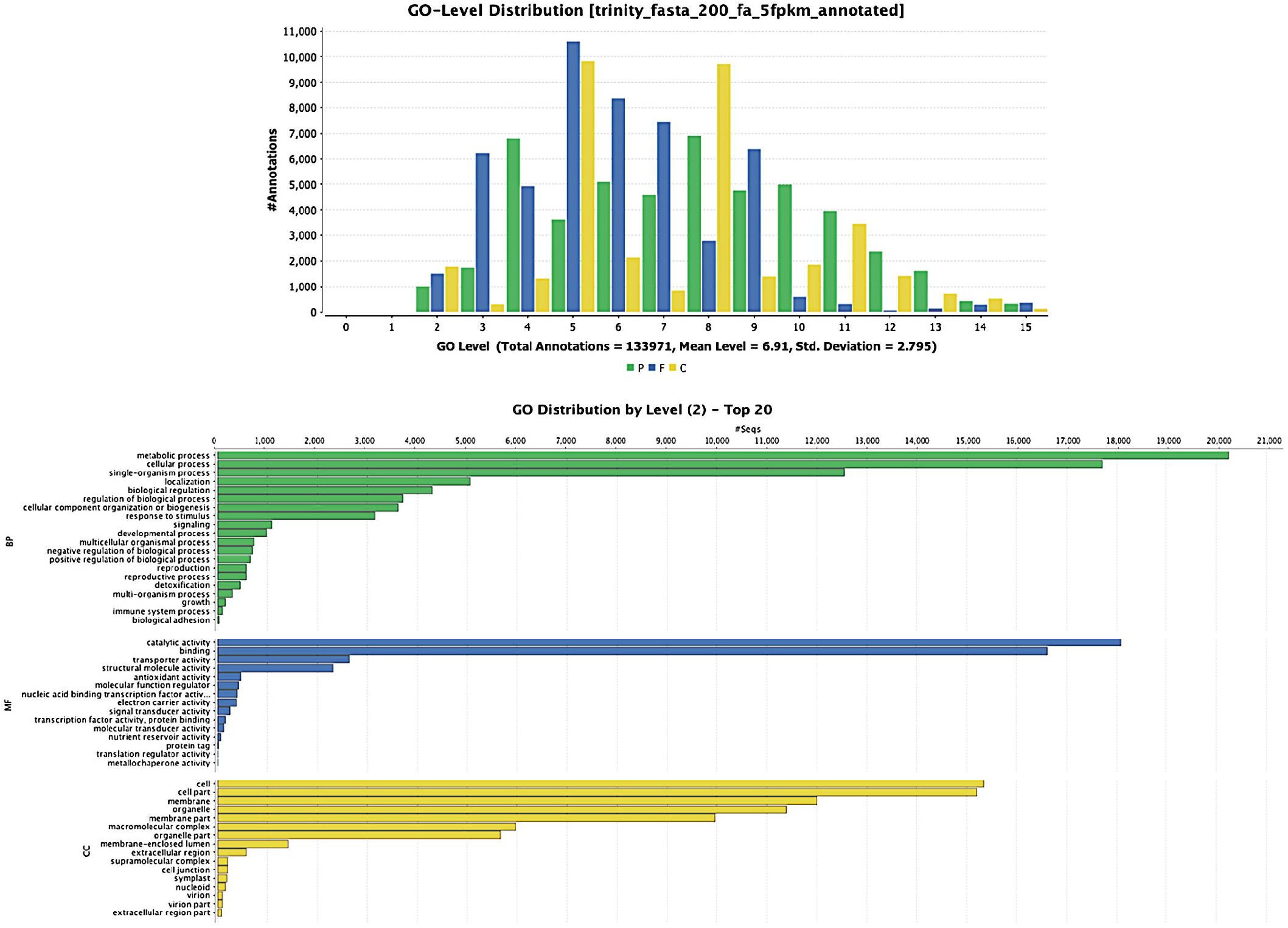
GO-level distributions in *S. bigelovii* transcriptome. **(A)** P, F and C represent the biological process (BP), molecular function (MF), and cellular component (CC), respectively. Total Annotations = 133,971, Mean Level = 6.91, and **(B)** Classification of *S. bigelovii* transcripts into functional categories (BP, MF, and CC) according to GO-terms on the basis of GO tool.

The pooled set of *S. bigelovii* transcripts were mapped against the InterPro database (http://www.ebi.ac.uk/interpro/) and assigned to 39,882 (21.4%) transcripts associated with 4,281 unique InterPro Families (Figure 4). The highest represented families were triphosphate hydrolase (1,089), protein kinase-like (918), and the alpha/beta hydrolase (ABH) superfamily (422) domains followed by the MFS transporter superfamily (387) and WD40/YVTN (410) domains. In addition, we identified transcripts involved in sugar transport (133), aquaporin transporter processing families (32), membrane transportation families (22), potassium transporter (21), Mg^2+^ transporter protein (12), and Na^+^/Ca^+2^ exchanger membrane region (7).

**Figure 4.**
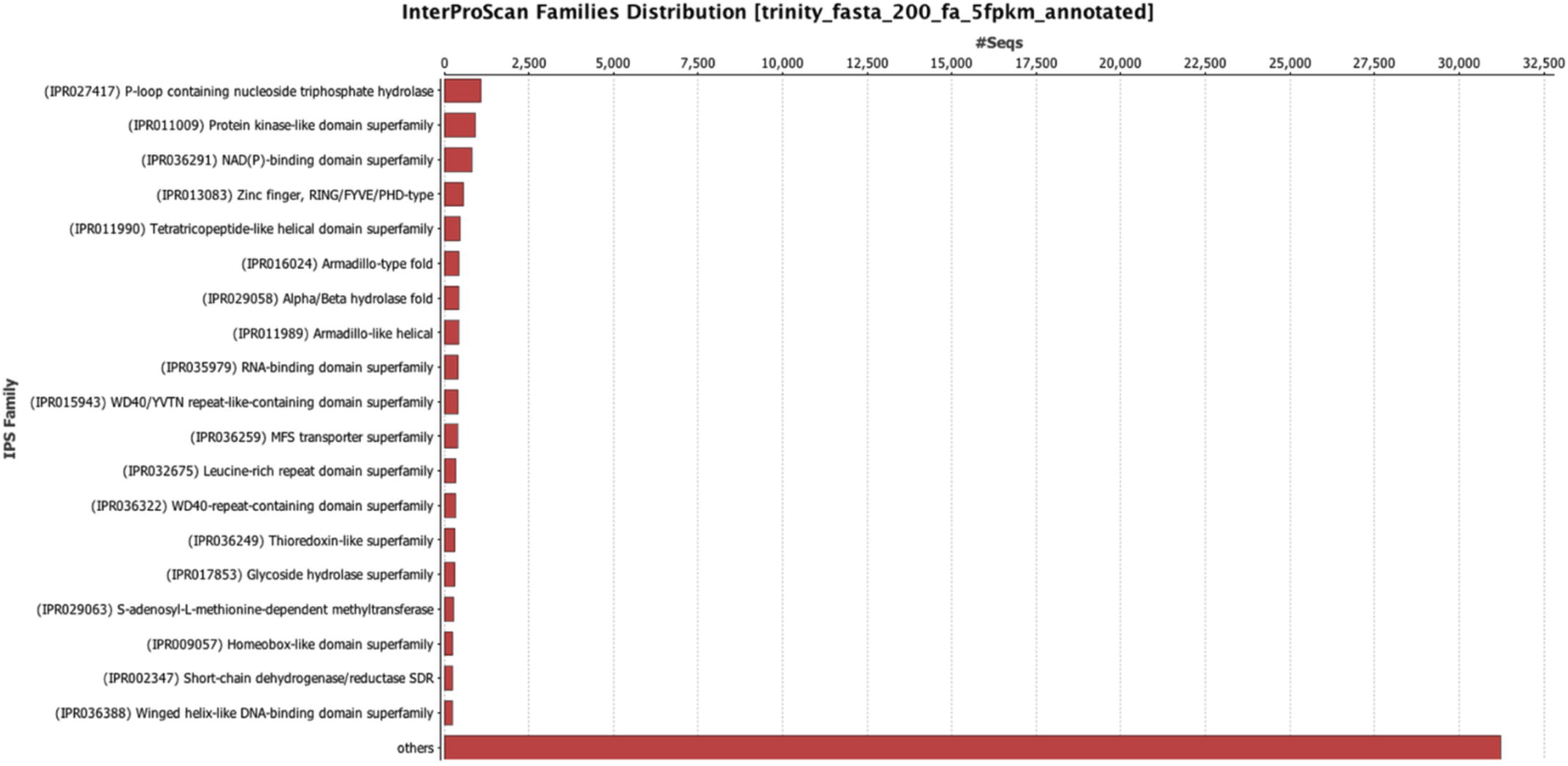
Distribution of protein domains predicted in the *S. bigelovii* transcriptome. Histogram of the 20 most abundant InterPro domains revealed by the InterProScan (IPS) annotation of assembled transcripts.

### *S. bigelovii* transcript expression analysis

For the differential expression analysis, the assembled transcript dataset identified as plant transcripts (49,914 transcripts) was used as an annotation reference when comparing transcripts from different *S. bigelovii* tissues. Differentially expressed transcripts (DETs) in roots, shoots, flowers, and seeds were identified using DESeqv1.8 for each tissue pairwise comparison. Transcripts were considered significantly differentially expressed based on a FDR of 5% and log2 |FC| > 2. Also, pairwise differential expression analysis of aerial plant tissues (shoots, seeds and flowers) transcripts compared to roots transcripts revealed up to 13,245 DETs depending on the comparison. The comparison between seeds and roots exhibited the highest number of DETs.

Hierarchical clustering of transcript abundance values of DETs between the four plant tissues shows the patterns of tissue-specific gene expression profiles that are clustered into four groupings (Figure 5). Overall and as expected, shoot profiles are more similar to flower profiles relative to root profiles while seed tissue expression is the most distinct from the other tissues. Transcripts cluster into four major groups (Clusters 1-4, Figure 5) that reflect the patterns of expression tissue specificity. Transcripts belonging to Cluster 1 are significantly up-regulated in seed tissue relative to shoot, root, and flower tissues (Cluster 1; 16,100). Cluster 2 (10,889) is enriched with transcripts up-regulated in flower and shoot tissues relative to root and seed tissues. A large set of transcripts in Cluster 4 were highly up-regulated in root tissue (Cluster 4; 11,681) relative to shoot tissue. Cluster 3 (11,244) on the other hand, is enriched with DETs with no clear visual pattern of tissue specific up- or down-regulation specific to a given tissue.

**Figure 5.**
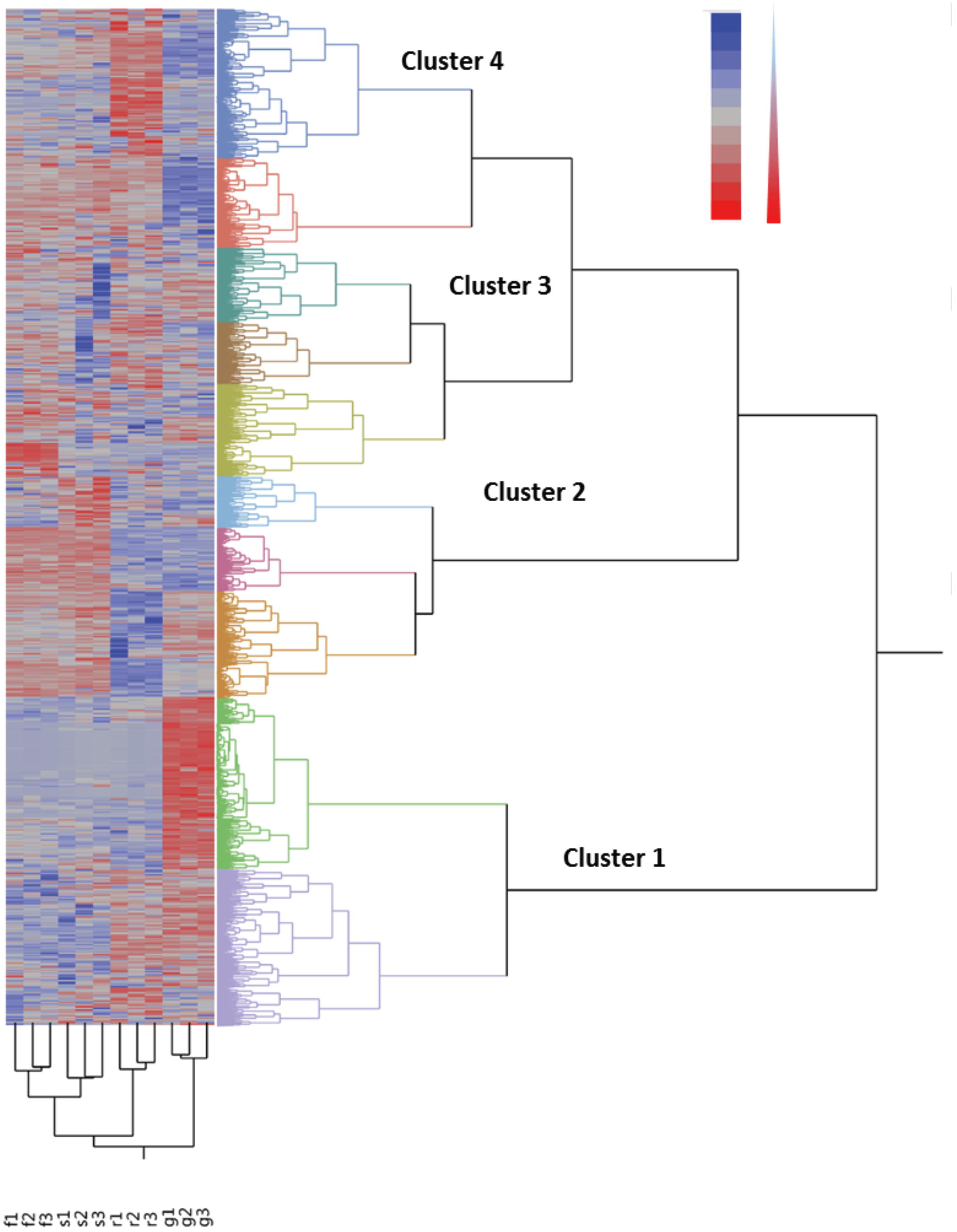
Global gene expression pattern of *S. bigelovii* transcriptome. Four major clusters of DET were identified in relation to tissue distribution. Heatmap displays global expression levels of DETs (rows) of all tissues based on FPKM values between all tissues. Columns represent samples (flower (F1, F2, F3), shoot (S1, S2, S3), root (R1, R2, R3), and seed (G1, G2, G3) tissues) while rows represent transcripts. For visualization purposes, the expression values were limited to 3 and −3.

A total of 3,362 transcripts were found commonly expressed in roots, shoots and seeds tissues while 1,261 were specifically differentially expressed in root and shoot, 1,101 specifically expressed in root and seed and 8,311 specifically differentially expressed in root and seed (Figure 6A). The number of DETs was highest between roots and aerial plant tissues (shoots, seeds and flowers) (Figure 6B). As expected, a relatively small number of DETs was identified between shoot and flower tissues with only 273 DETs down-regulated and 948 DETs up-regulated in shoot tissues.

**Figure 6.**
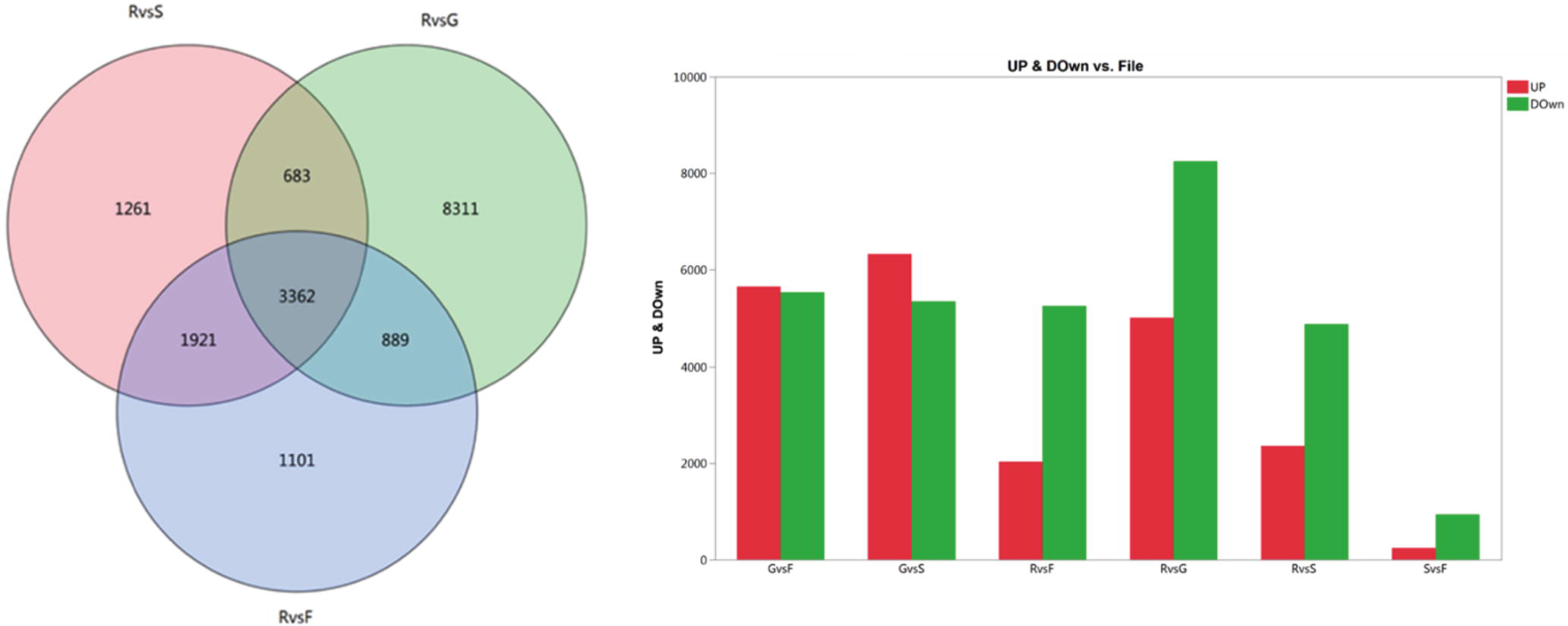
Global DETs comparison between all four tissue libraries. **(A)** Distribution of transcripts differentially expressed between roots and shoots (red), roots and seeds (green), roots and flowers (blue). Statistics were performed using the DESeqv1.8 methods with FDR 0.05 and FC=2. **(B)** The red columns indicate the up-regulated DETs and the green columns represent the down-regulated DETs in four pair-wise tissue comparisons.

A total of 7,273 transcripts were differentially expressed between root and flower tissue, of which 5,452 were down-regulated and 1,821 up-regulated in root tissue. We identified 13,245 DETs between root and seed tissues, with 8,586 being down-regulated and 4,659 being up-regulated in root tissue. Moreover, we identified 7,227 DETs between root and shoot tissues, of which 4,345 were down-regulated and 2,882 were up-regulated in root tissue (Figure 6B).

### Gene ontology analysis and identification of potential salt tolerance transcripts in shoots and roots

Previous work analyzed the global transcriptome profile in *Salicornia europaea* roots and shoots (Furtado et al., 2019). This work done in very different conditions that those experienced in Abu Dhabi. In order to understand the metabolic response of *S. bigelovii* to the extreme environmental stresses in Abu Dhabi, we performed an in-depth GO functional analysis of DETs between root and shoot tissues. All DETs were grouped into 48 GO classes belonging to 15 CC terms, 12 MF terms, and 21 BP terms, (Figure S4). Similar GO distribution was observed for up-regulated and down-regulated transcripts in root as compared to shoot tissues. For BPs, metabolic process and cellular processes were the highest differentially expressed terms; we also took notice of the high number of transcripts belonging to response to stimulus (410) upregulated in roots. For MF, the two most expressed transcripts were related to catalytic activity and binding with transporter activity while for CC, the highest differentially expressed terms were cell, cell part and organelle (Figure S4).

Among transcripts involved in transport activity (GO:0005215) and response to stimulus (GO:0050896), we identified essentially three aquaporin transcripts: two encoding tonoplast membrane aquaporins (TIP) (TRINITY_DN208648, TRINITY_DN200599) and one encoding plasma membrane aquaporin (PIP) (TRINITY_DN209082, K09872) that were up-regulated in root as compared to shoot tissues (FDR < 5%; log2 FC = 8.43 and FDR < 5%; log2 FC = 2.54, respectively) (Figure 7).

**Figure 7.**
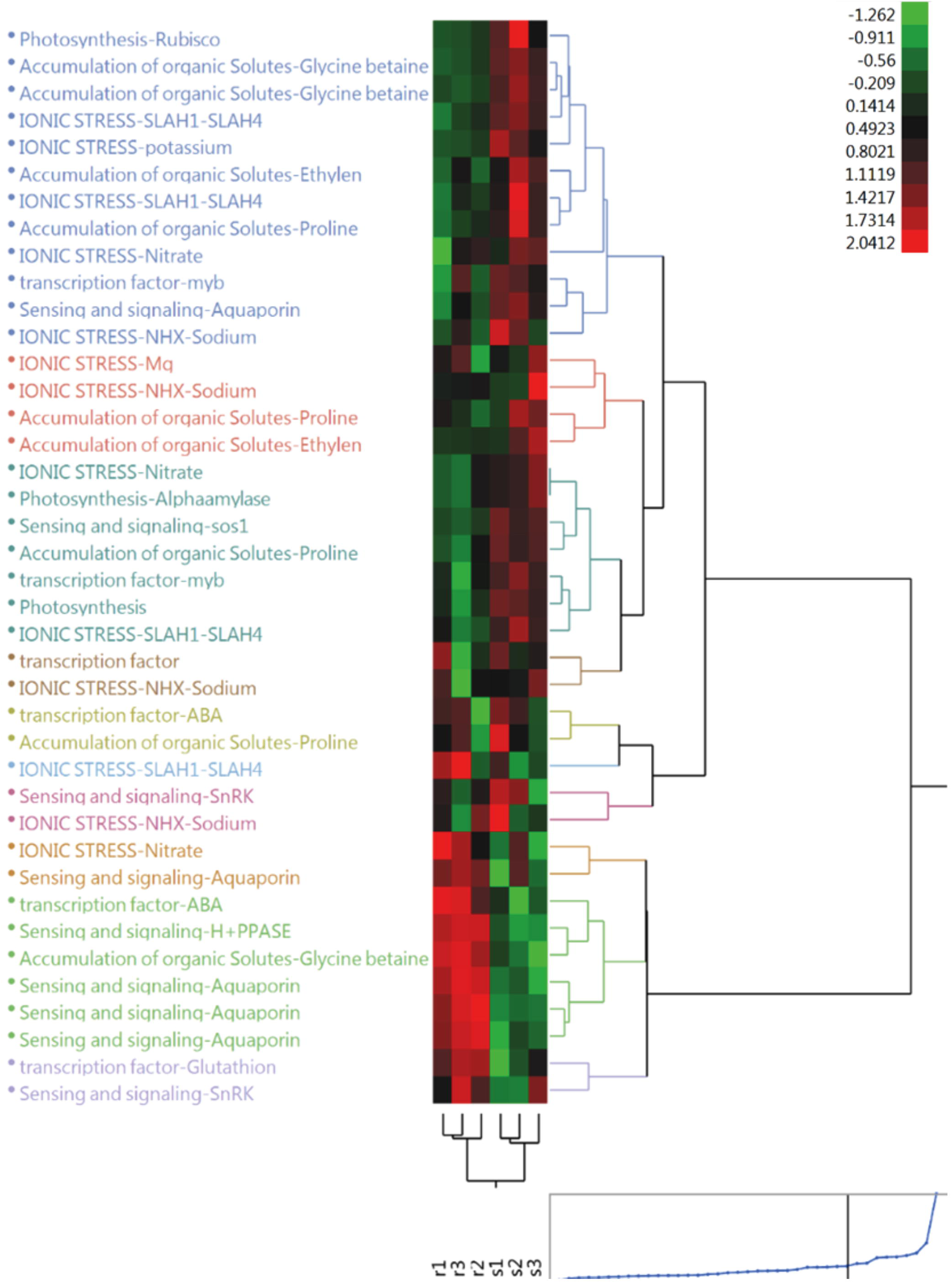
Heatmap of transcripts implicated in salt stress tolerance based on FPKM units of DETs between shoots (S1, S2, S3) and roots (R1, R2, R3).

Transcripts encoding vacuolar H^+^-pyrophosphatase (H^+^-PPase) (TRINITY_DN206459-K23025) were also upregulated in root tissue during sea effluent water irrigation. However, transcripts associated with salt oversensitive stress (SOS1) (TRINITY_DN220726) exhibited higher expression in shoot tissue when compared to the roots.

A considerable number of DETs (GO:0005215; 398 transcripts) encoding for a diversity of ion transporters such as sodium antiporter, calcium and potassium transporter were found to be up-regulated in shoot relative to root tissues. The predicted antiporter NHX4 (TRINITY_DN207378) and the transcript encoding for the high affinity potassium transporter, HKT6 (TRINITY_DN119119) were highly upregulated in shoots compared to roots (FDR < 5%; log2 FC = 3.82 and FDR < 5%; log2 FC = 6.48, respectively). The Na^+^/H^+^ antiporter NHX1 (TRINITY_DN224256), the predicted antiporter NHX2 (TRINITY_DN221143) and the Ca^2+^ transporting ATPase (TRINITY_DN110967) are also up-regulated in shoots (FDR < 5%; log2 FC = 2.82 and FDR < 5%; log2 FC = 2.79, respectively).

Several DETs encoding for proteins involved in the synthesis and metabolism of osmolytes such as proline and glycine betaine were identified. We highlight in particular two transcripts encoding for two enzymes in the glycine betaine pathway: the betaine aldehyde dehydrogenase (BADH) (GO:0008152, TRINITY_DN226919, FDR < 5%; log2 FC = 2.31) and the choline monooxygenase (CMO) (TRINITY_DN213512, FDR < 5%; log2 FC = 2.01) upregulated in shoots relative to roots. Interestingly, another transcript encoding for BADH (TRINITY_DN231325) was highly upregulated in root tissues (FDR < 5%; log2 FC = 3.59). In addition, we found transcripts mapping to the proline metabolism pathway (IPR0 18800). We identified the Δ-pyrroline-5-carboxylate synthetase (P5CS) (TRINITY_DN223373) transcript and the proline-rich protein (PRP) (TRINITY_DN218724) to be upregulated in shoots (FDR < 5%; log2 FC = 2.07, FDR < 5%; log2 FC = 2.44, respectively).

Transcription factors (TFs) play vital roles in regulating plant resistance mechanisms under abiotic stress. Based on the GO term (GO: 0001071), 175 DETs were identified as nucleic acid binding and TFs with the majority of the transcripts being up-regulated in root tissue. The most abundant classes of up-regulated TFs in roots included WRKY (TRINITY_DN192570, TRINITY_DN221294) (FDR < 5%; log2 FC = 7.1 and FDR < 5%; log2 FC = 7.99, respectively) while MYB (TRINITY_DN155341, TRINITY_DN190623) (FDR < 5%; log2 FC = 3.92 and FDR < 5%; log2 FC = 3.32, respectively) TFs were up-regulated in shoots.

### Evaluation and validation of selected genes using qRT-PCR

We performed qRT-PCR to validate the relative changes in transcript levels observed in the transcriptome data. Nine genes randomly selected from transcriptome reads exhibiting differential expression patterns in the *de novo* assembly analysis were compared with the data obtained by qRT-PCR (Figure 8). The qRT-PCR data revealed similar relative expression trends between the selected *de novo* assembly transcripts. For example, the SOS1, the P5PR, and NHX1 genes displayed the same relative levels of expression in both the qRT-PCR and transcriptome data. These genes were up-regulated in the shoot as compared to root tissues. The aquaporins (AQPs), H^+^-PPase, and glycine betaine genes also showed the same relative expression patterns in both the qRT-PCR and *de novo* transcriptome data: they were down-regulated in shoot relative to root tissues. Overall, we observed some variability in DETs between the qRT-PCR data and the *de novo* transcriptome data.

**Figure 8.**
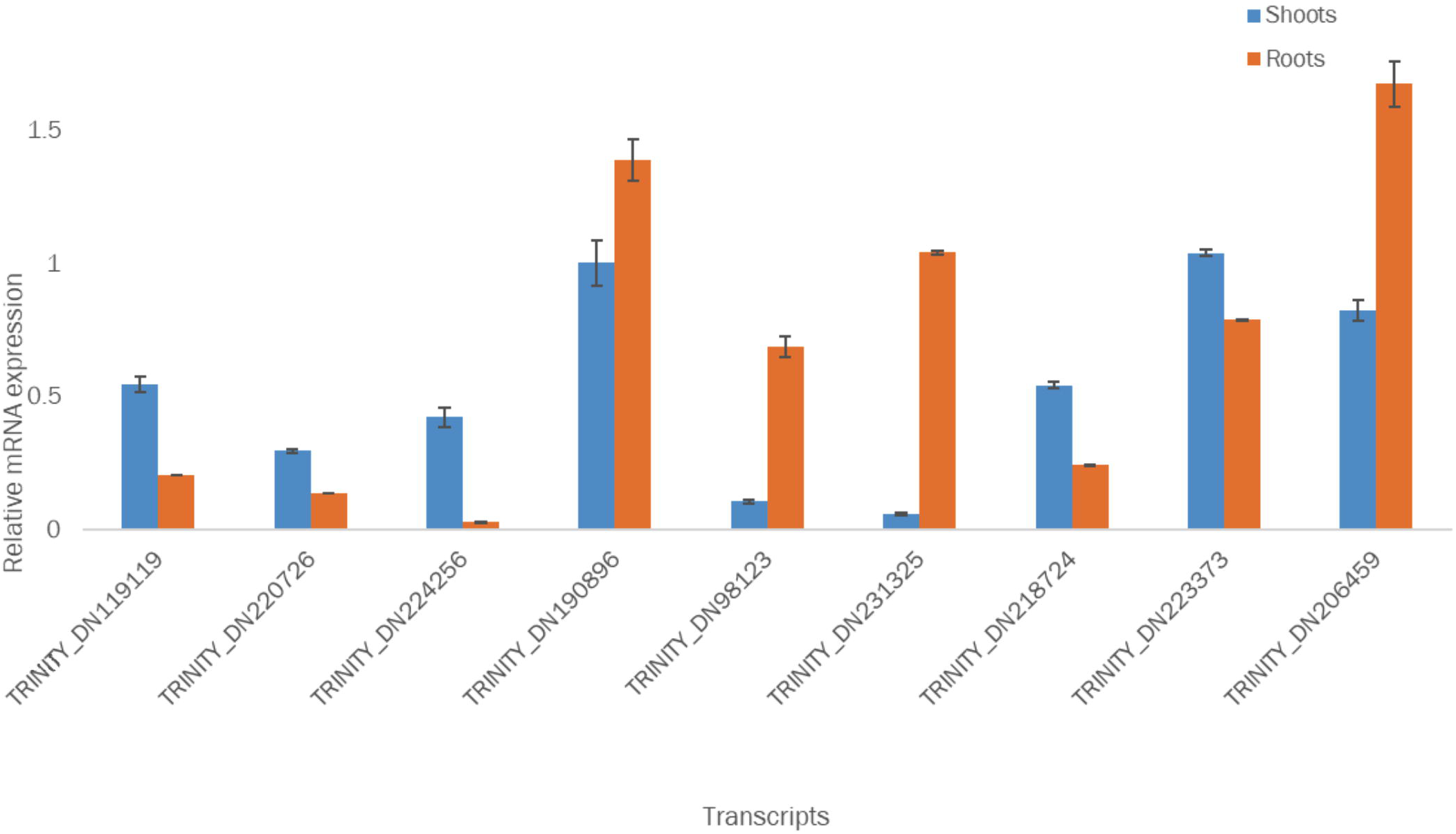
The expression validation of candidate transcripts implicated in salt stress tolerance in *S. bigelovii* by qRT-PCR. Error bars represent the mean (± SD) of three replicates.

## Discussion

Soil salinity is the primary environmental factor affecting agriculture and crop productivity (Flowers and Colmer, 2008; Zörb et al., 2019). The lack of water along with a rise in thermal stress are anticipated to worsen, thus further reducing agriculture productivity. Halophyte plant crop plants offer an opportunity to diversify and augment agriculture productivity (Panta et al., 2014). Unfortunately, our understanding of the global molecular mechanisms underlying high salt tolerance in halophyte plants under environmental stress is limited in part due to scarcity of genomic and transcriptional studies. The true halophyte plant *S. bigelovii* exhibits high salt tolerance typical of its natural environment (Kadereit et al., 2007). This plant offers a unique opportunity to study the global transcriptional response to high salt stress in salt tolerant plants.

One barrier to understanding the global plant response to its environment is the lack of transcriptional data from multiple tissues as they respond to environmental stresses over the course of plant growth and maturation cycle.

Furtado et al. investigated the DETs between shoots and roots of *S. europaea* in at two sites, Spa Park (site 1) and Ciechocinek (site 2) in central Poland where the environmental conditions are much different than those experienced in Abi Dhabi (Furtado et al., 2019). The average monthly temperature was between 1.5°C and 22.1°C at site 1 and −2.6°C and 17.8°C at site 2 in January and August respectively whereas the average monthly temperature in Abu Dhabi ranged from 23.9°C in January to 41.1°C in August (Statistics Center, 2018). The annual precipitation was 379.4 mm at site 1 and 680.2 mm at site 2, as compare to 23 mm in Abu Dhabi. Site salinities were much lower at their sites (9.2 ppt to 21.5 ppt and 7.4 ppt and 11.8 ppt in spring and fall) than those in Abu Dhabi (34.6 ppt and 42.6 ppt in March and August).

*S. bigelovii* plants grown at the SEAS pilot facility were exposed to the annual environmental conditions, such as temperature, humidity, salinity and length of day, between the Abu Dhabi winter and late summer seasons. These conditions represent varying physical stresses experienced by plants through the stages of growth and maturation.

We have used RNA sequencing and performed a transcriptome analysis to study the global gene expression profiles in four tissues of *S. bigelovii* under saline environmental conditions. To our knowledge, this represents the first global transcriptome data available for *S. bigelovii*.

### *De novo* assembly and analysis

One major advantage of RNA-Seq analysis is its capacity to identify previously unknown transcripts through *de novo* assembly. We identified a total of 66,943 transcripts after *de novo* assembly, of which 72.63% were successfully annotated in the GO database. The results show a total of 18,321 transcripts classified as “Others” (27.38%) with no matches to known transcripts. These unknown transcripts may represent novel *S. bigelovii* transcripts some of which might be involved in the mechanisms of salt tolerance.

The BLAST results show that *B. vulgaris maritima* had the highest number of identified transcripts. This result is as expected as *B. vulgaris maritima* is closely related to *S. bigelovii*. The next species identified by BLAST is *S. oleracea*. These two species accounts for more than 58% of all top-species hits. Additionally, our analysis indicated the presence of transcripts of non-plant origin, mainly coming from bacteria and fungi.

We expected to identify non-plant transcripts as plants colonized by endophytes often display increased tolerance to abiotic stresses such as salinity and drought (Singh et al., 2011). It has been shown that endophytes actively colonize plants, interact with their host, and frequently show beneficial effects on plant growth and health (Vaishnav et al., 2019). Still, the mechanisms of plant-endophyte interaction and fungal adaption to the plant environment are poorly understood. These data allude to the identity and metabolic processes between plants and endophytes.

After removal of transcripts belonging to non-plant organisms, we were left with 49,914 transcripts for the analysis of *S. bigelovii* DETs. Our data provides the first resource for global gene identification and regulation analysis in *S. bigelovii* and other strict halophytes.

### Global differentially expressed transcripts analysis

Plants subjected to salt stress display complex metabolic interactions between signal transduction networks, transcriptional regulation, and stress gene expression (Deinlein et al., 2014). We identified a set of DETs that were significantly upregulated in the above ground plant tissues (shoot, seed, and flower) relative to the underground tissue (root) (Figure 6). The analysis identified several key stress response DETs in specific tissues that were not previously reported in the literature. We focused on comparing DETs between shoot and root tissues to identify genes known to be associated with salt tolerance in the halophyte *S. bigelovii*. Using qRT-PCR, the RNA-Seq results for 9 randomly selected salt stress-associated genes were validated. The qRT-PCR results agreed with the overall differential expression trends observed in the transcriptome analysis. Future characterization of the role of specific transcripts is required to develop a better understanding of the relationship between environmental stresses and gene expression and regulation profiles in *S. bigelovii*.

### Growth and ion balance under agricultural effluent water

Salinity and drought stress reduces water transport rates across membranes (Osakabe et al., 2014). The uptake of water by plants is highly dependent on regulation of AQPs (Maurel et al., 2015). AQPs are transmembrane proteins that facilitate uptake of soil water and regulate root hydraulic conductivity. They are also involved in cellular compartmentalization of water and are thought to play a role in maintaining osmosis and turgor of plant cells in halophytes (Berger et al., 2010).

There is still much debate regarding salt dependent regulation of AQPs. For instance, even though *Kochia sieversiana* can subsist at high salinity, most AQP genes were significantly up-regulated in low, but not high salinity stressed roots (Zhao et al., 2017). In contrast, the halophyte *Schrenkiella parvula* expressed a high number of AQPs for tolerance to salt toxicity with TIP2 being highly expressed in *S. parvula* root as compared to shoot tissues (Loqué et al., 2005; Oh et al., 2014). Our analysis shows that TIPs (TIP1 and TIP2) were significantly up-regulated in roots of *S. bigelovii* when compared to previous halophyte literature (Salazar, 2017). This presents tantalizing data that suggest multiple pathways for regulation of AQPs in halophytes.

Cellular ion transport across the tonoplast into vacuoles is maintained by the proton motive force (PMF) generated by the vacuolar H^+^-PPase (Gaxiola et al., 2007). It has been reported that both the H^+^-PPase and V-ATPase transport activity in *S. bigelovii* (Ayala et al., 1996) increased upon the addition of NaCl to the growth medium. We observed that *Sb*H^+^-PPase is also upregulated in root as compared to shoot tissues.

Up-regulation of AQPs and H^+^-PPases is counterbalanced by high expression of several monovalent ion transporters. These ion transporters regulate Na^+^, K^+^, and Cl^-^ transport, which are necessary for increased salt tolerance. The ion transport counterbalance is undertaken by vacuolar membrane Na^+^/H^+^ exchangers (NHX) and is driven by the intracellular electrochemical gradient of protons membrane, or NHX1 in the tonoplast. We identified three *Sb*NHX genes homologous to NHX genes found in *S. oleracea, B. vulgaris* and *S. europaea* (Barkla et al., 1995; Su et al., 2003). The upregulation of the NHX1 (TRINITY_DN224256) transcript in shoot tissue was confirmed by qRT-PCR analysis. This was previously shown in the halophyte *Mesembryanthemum crystallinum* and NHX1 expression is thought to enhance acclimation to increasing environmental salinity (Barkla et al., 1995; Su et al., 2003). These studies show that vacuolar Na^+^/H^+^ antiporters are important for increasing plant salt tolerance through Na^+^ sequestration (Su et al., 2003).

The HKT6 gene is a subfamily member of low-affinity Na^+^/K^+^ transporters (Platten et al., 2006). We identified only one high affinity K^+^ transporter transcript that is homologous to the HKT6 gene in *B. vulgaris* and it was highly up-regulated in *S. bigelovii* shoot relative to root tissues. In line with an observation in *Salicornia dolichostachya*, where the *Arabidopsis thaliana* HKT1 orthologous transcript was not detectable in root tissue transcriptome data (Katschnig et al., 2015), we are also unable to detect HKT1 expression in both shoot and root tissue. Low HKT1 transcript expression in roots has been observed in other species that are members of the Amaranthaceae: *M. crystallinum* (Su et al., 2003) and *Suaeda salsa* (Shao et al., 2008). These differences in HKT1 expression levels have been attributed to root tissue Na^+^ ‘accumulation strategy’ by these species.

The efflux of Na^+^ across the plasma membrane is regulated by the SOS1 Na^+^/H^+^ antiporter (Shi et al., 2000). SOS1 mediates Na^+^ efflux to the apoplast against the electrochemical potential via secondary active transport (Ji et al., 2013). Researchers have suggested that *Ss*SOS1 may mediate Na^+^ efflux in leaves and roots but reduce Na^+^ through long distance transfer regulation in stems minimizing Na^+^ toxicity and maintaining homeostasis during salt stress (Song and Wang, 2015). In the halophyte *S. salsa*, the expression of the *Ss*SOS1 in roots, stems and leaves is induced by salt stress (Wang et al., 2013). In *T. halophila*, there was a sevenfold increase of *Th*SOS1 transcript expression levels in root relative to shoot tissues (Katschnig et al., 2015). Interestingly, in our current study, expression of the *Sb*SOS1 is up-regulated in shoots rather than in root tissue. In *S. dolichostachya* shoot tissue, high expression of *Sd*SOS1 shows complete suppression of *Sd*HKT1 (Katschnig et al., 2015).

In conclusion, in *S. bigelovii* the *Sb*SOS1, *Sb*NHX, and *Sb*HKT6 genes are up-regulated in shoots, while aquaporins are up regulated in roots. These data present one more instance in which the regulation of salt stress transcripts differs in *S. bigelovii* relative to what was observed in other halophyte plant species tissues. Together with the previous observations, our findings suggest that *S. bigelovii* is a salt accumulating species, but the salt accumulation in its shoots occurs through an unknown mechanism (Salazar, 2017; Furtado et al., 2019).

### Compatible solutes and osmolyte production

Plants generally compartmentalize Na^+^ into vacuoles in order to avoid Na^+^ toxicity. To combat the osmotic stress caused by higher concentrations of Na^+^ in the vacuoles, plants accumulate organic compatible solutes and osmolytes, such as betaine and proline, in their cytoplasm (Parida and Das, 2005; Munns and Tester, 2008).

We identified the Δ-pyrroline-5-carboxylate synthetase (P5CS, TRINITY_DN223373) transcript which has been shown to be involved in proline biosynthesis in plants. The synthesis and transport of these amino acids promote salt tolerance in most plants (Hasegawa et al., 2000; Munns, 2002; Flowers and Colmer, 2008; Munns and Tester, 2008). In the euhalophyte, *S. salsa*, P5CS was upregulated by salinity in different tissues (Wang et al., 2002). The proline-rich protein (*PRP*) homologue gene in *B. vulgaris* was also expressed is *S. bigelovii* shoot and root tissues. It is highly up-regulated in the *S. bigelovii* shoot tissue and down-regulated in root tissue. These findings suggest that the halophyte *S. bigelovii* synthesizes and transports various amino acids to maintain cell turgor pressure under osmotic stress.

Betaine aldehyde dehydrogenase (BADH) is a key enzyme for glycine betaine synthesis, which plays an important role in improving plant tolerance to salinity (Fitzgerald et al., 2009). We have identified two BADH transcripts, one upregulated in shoots (TRINITY_DN213512) and the other in roots (TRINITY_DN231325). Further analyses are needed to measure the levels of these osmolytes and amino acids to confirm their cellular concentration.

The regulation of osmolytes and metabolites production is also maintained by TFs. They play a significant role in plant development, reproduction, intercellular signaling, and cell cycle (Singh et al., 2002). The WRKY and MYB TFs are unique to plants (Riechmann et al., 2000). Several studies link specific members of WRKY and MYB TF families to plant stress responses (Rushton et al., 2010; Li et al., 2019; Tang et al., 2019). We identified transcripts coding for 2 WRKYTFs; WRKY65 and WRKY75. These are present in *B. vulgaris* and are up-regulated in *S. bigelovii* shoots. This TF family has also been associated with plant stress responses to anaerobic stress and regulation of secondary metabolite during stress conditions induced by the presence of pathogens (Phukan et al., 2016). In *Arabidopsis*, WRKY75 initiates stress response by regulating nuclear encoded organelle proteins (Van Aken et al., 2013). Several TF MYB proteins, such as the MYB44 (TRINITY_DN219454) and MYB28 (TRINITY_DN190623) are up-regulated in *S. bigelovii* roots. Overexpression of *Sb*MYB44 enhanced the growth of yeast cells under both ionic and osmotic stresses (Shukla et al., 2015). Overall, salt tolerance in halophytes is a convoluted network that requires highly regulated and coordinated responses of genes and metabolites.

*S. bigelovii* is a valuable halophyte plant adapted to growth in coastal deserts with potential use as a food crop. Even though studies show the beneficial nutritional and health properties of *Salicornia* species for use as food and fodder, we lack information regarding its genetic makeup, metabolic potential, and salt tolerance mechanisms. Understanding salt tolerance in plants at the whole plant, organelle, and molecular level can point to the selection of salt tolerant food crop genotypes that have increased crop productivity and quality.

We used RNA sequencing data to *de novo* assemble the global transcriptome for *S. bigelovii* grown under Abu Dhabi’s desert environmental conditions. Transcriptomic data from four tissues were analyzed with a special emphasis on identifying tissue-specific expression patterns previously implicated in salt stress response. The transcriptome results were validated by gene expression analysis using qRT-PCR. The identification of *S. bigelovii* specific transcripts can be exploited for elucidating metabolic systems, osmotic stress related secondary metabolite production, and oil biosynthesis in *S. bigelovii*.

It comes as no surprise that the tissue specific expression levels of genes between *S. bigelovii* and other plants vary and are dependent on the cellular function in response to abiotic stress. These data also suggest that TFs are a key member of these events as our data suggest they are involved in regulation key cellular metabolic processes in plants. To summarize, the results of this study provide the first transcriptome sequencing of the strict halophyte, *S. bigelovii*.

## Data Availability Statement

All RNA sequencing data have been deposited at the NCBI in the Short Read Archive database (Bioproject ID: PRJNA607385, Biosample ID: SUB6825665) and is available under request.

## Author Contributions

HC designed the study, performed all field experiments, collection and preparation of tissue samples, RNA isolation, qPCR-based expression analysis for transcriptomics study, performed RNA-Seq validations, measurement of gene expression from qPCR, and wrote and revised the manuscript. MV performed the bioinformatics analyses including genome assembly, annotation, and RNA-Seq analysis. MD prepared RNA-Seq libraries and performed high throughput sequencing. YI provided critical inputs for RNA-Seq data analysis and presentation. AH critically reviewed the manuscript. HHH and revised, reviewed, submitted the manuscript, and provided the resources All authors have read and approved the final manuscript.

## Acknowledgments

We would like to thank all the people at the Seawater Energy and Agriculture System pilot facility for their help in sample collection and Dr. Runyararo M. Nyadzayo for all her constructive comments and edits to the manuscript.

## Supplementary Material

The Supplementary Material for this article can be found online.

## References

Ahmad, R., Jamil, S., Shahzad, M., Zörb, C., Irshad, U., Khan, N., et al. (2017). Metabolic Profiling to Elucidate Genetic Elements Due to Salt Stress. CLEAN – Soil, Air, Water 45(12), 1600574. doi:10.1002/clen.201600574.

Andrews, S. (2010). “FASTQC. A quality control tool for high throughput sequence data”.).

Anwar, F., Bhanger, M.I., Nasir, M.K., and Ismail, S. (2002). Analytical characterization of *Salicornia bigelovii* seed oil cultivated in Pakistan. J Agric Food Chem 50(15), 4210–4214. doi:10.1021/jf0114132.

Ashburner, M., Ball, C.A., Blake, J.A., Botstein, D., Butler, H., Cherry, J.M., et al. (2000). Gene Ontology: tool for the unification of biology. Nature Genetics 25(1), 25–29. doi:10.1038/75556.

Atkinson, N.J., and Urwin, P.E. (2012). The interaction of plant biotic and abiotic stresses: from genes to the field. Journal of experimental botany 63(10), 3523–3543.

Ayala, F., O’Leary, J.W., and Schumaker, K.S. (1996). Increased vacuolar and plasma membrane H^+^-ATPase activities in *Salicornia bigelovii* Torr. in response to NaCl. Journal of Experimental Botany 47(1), 25–32. doi:10.1093/jxb/47.1.25.

Barkla, B.J., Zingarelli, L., Blumwald, E., and Smith, J. (1995). Tonoplast Na^+^/H^+^ antiport activity and its energization by the vacuolar H^+^-ATPase in the halophytic plant *Mesembryanthemum crystallinum* L. Plant Physiol 109(2), 549–556. doi:10.1104/pp.109.2.549.

Berger, B., Parent, B., and Tester, M. (2010). High-throughput shoot imaging to study drought responses. J Exp Bot 61(13), 3519–3528. doi:10.1093/jxb/erq201.

Bolger, A.M., Lohse, M., and Usadel, B. (2014). Trimmomatic: a flexible trimmer for Illumina sequence data. Bioinformatics 30(15), 2114–2120. doi:10.1093/bioinformatics/btu170.

Boyer, J.S. (1982). Plant productivity and environment. Science 218(4571), 443–448. doi:10.1126/science.218.4571.443.

Bray, R.A., Bailey-Serres, J., and Weretilnyk, E. (2000). “Response to abiotic stress,” in Biochemistry and molecular biology of plants, eds. B.B. B., G. W. & J.R. L. (New York, NY USA: American Society of Plant Physiologists), 1158–1203.

Consortium, T.U. (2018). UniProt: a worldwide hub of protein knowledge. Nucleic Acids Research 47(D1), D506–D515. doi:10.1093/nar/gky1049.

Deinlein, U., Stephan, A.B., Horie, T., Luo, W., Xu, G., and Schroeder, J.I. (2014). Plant salt-tolerance mechanisms. Trends in plant science 19(6), 371–379. doi:10.1016/j.tplants.2014.02.001.

El-Mallah, M.H., Turui, T., and El-Shami, S. (1994). Detailed studies on seed oil of *Salicornia* SOS-7 cultivated at the egyptian border of Red Sea. Grasas Y Aceites.

Fan, P., Nie, L., Jiang, P., Feng, J., Lv, S., Chen, X., et al. (2013). Transcriptome analysis of *Salicornia europaea* under saline conditions revealed the adaptive primary metabolic pathways as early events to facilitate salt adaptation. PLoS One 8(11).

Fitzgerald, T.L., Waters, D.L., and Henry, R.J. (2009). Betaine aldehyde dehydrogenase in plants. Plant biology 11(2), 119–130.

Flowers, T.J., and Colmer, T.D. (2008). Salinity tolerance in halophytes. New Phytologist, 945–963.

Flowers, T.J., Galal, H.K., and Bromham, L. (2010). Evolution of halophytes: multiple origins of salt tolerance in land plants. Functional Plant Biology 37(7), 604–612.

Flowers, T.J., Munns, R., and Colmer, T.D. (2015). Sodium chloride toxicity and the cellular basis of salt tolerance in halophytes. Annals of botany 115(3), 419–431.

Furtado, B.U., Nagy, I., Asp, T., Tyburski, J., Skorupa, M., Golebiewski, M., et al. (2019). Transcriptome profiling and environmental linkage to salinity across *Salicornia europaea* vegetation. BMC plant biology 19(1), 427.

Gaxiola, R.A., Palmgren, M.G., and Schumacher, K. (2007). Plant proton pumps. FEBS letters581(12), 2204–2214.

Ghosh, P., Westhoff, P., and Debnath, D. (2019). “Biofuels, food security, and sustainability,” in Biofuels, Bioenergy and Food Security, eds. D. Debnath & S.C. Babu. Academic Press), 211–229.

Gill, S.S., and Tuteja, N. (2010). Reactive oxygen species and antioxidant machinery in abiotic stress tolerance in crop plants. Plant physiology and biochemistry 48(12), 909–930.

Glenn, E., Miyamoto, S., Moore, D., Brown, J.J., Thompson, T.L., and Brown, P. (1997). Water requirements for cultivating *Salicornia bigelovii* Torr. with seawater on sand in a coastal desert environment. Journal of Arid Environments 36(4), 711–730.

Glenn, E.P., Anday, T., Chaturvedi, R., Martinez-Garcia, R., Pearlstein, S., Soliz, D., et al. (2013). Three halophytes for saline-water agriculture: An oilseed, a forage and a grain crop. Environmental and Experimental Botany 92, 110–121.

Glenn, E.P., Brown, J.J., and Blumwald, E. (1999). Salt tolerance and crop potential of halophytes. Critical reviews in plant sciences 18(2), 227–255.

Glenn, E.P., Coates, W.E., Riley, J.J., Kuehl, R.O., and Swingle, R.S. (1992). *Salicornia bigelovii* Torr.: a seawater-irrigated forage for goats. Animal Feed Science and Technology 40(1), 21–30.

Glenn, E.P., Jed Brown, J., and O’Leary, J.W. (1998). Irrigating crops with seawater. SCIENTIFIC AMERICAN-AMERICAN EDITION- 279, 76–81.

Gomes-Filho, E., Lima, C.R.F.M., Costa, J.H., da Silva, A.C.M., da Guia Silva Lima, M., de Lacerda, C.F., et al. (2008). Cowpea ribonuclease: properties and effect of NaCl-salinity on its activation during seed germination and seedling establishment. Plant Cell Reports 27(1), 147–157. doi:10.1007/s00299-007-0433-5.

Götz, S., García-Gómez, J.M., Terol, J., Williams, T.D., Nagaraj, S.H., Nueda, M.J., et al. (2008). High-throughput functional annotation and data mining with the Blast2GO suite. Nucleic acids research 36(10), 3420–3435.

Grabherr, M.G., Haas, B.J., Yassour, M., Levin, J.Z., Thompson, D.A., Amit, I., et al. (2011). Full-length transcriptome assembly from RNA-Seq data without a reference genome. Nature biotechnology 29(7), 644.

Hao, X., Horvath, D.P., Chao, W.S., Yang, Y., Wang, X., and Xiao, B. (2014). Identification and evaluation of reliable reference genes for quantitative real-time PCR analysis in tea plant (*Camellia sinensis* (L.) O. Kuntze). International journal of molecular sciences 15(12), 22155–22172. doi:10.3390/ijms151222155.

Hasegawa, P.M., Bressan, R.A., Zhu, J.-K., and Bohnert, H.J. (2000). Plant cellular and molecular responses to high salinity. Annual review of plant biology 51(1), 463–499.

Huerta-Cepas, J., Szklarczyk, D., Forslund, K., Cook, H., Heller, D., Walter, M.C., et al. (2016). eggNOG 4.5: a hierarchical orthology framework with improved functional annotations for eukaryotic, prokaryotic and viral sequences. Nucleic acids research 44(D1), D286–D293.

Jenks, M.A., Hasegawa, P.M., and Jain, S.M. (2007). Advances in molecular breeding toward drought and salt tolerant crops. Springer.

Ji, H., Pardo, J.M., Batelli, G., Van Oosten, M.J., Bressan, R.A., and Li, X. (2013). The Salt Overly Sensitive (SOS) pathway: established and emerging roles. Molecular plant 6(2), 275–286.

Kadereit, G., Ball, P., Beer, S., Mucina, L., Sokoloff, D., Teege, P., et al. (2007). A taxonomic nightmare comes true: phylogeny and biogeography of glassworts (*Salicornia* L., *Chenopodiaceae*). Taxon 56(4), 1143–1170.

Kadereit, G., Mucina, L., and Freitag, H. (2006). Phylogeny of *Salicornioideae (Chenopodiaceae*): diversification, biogeography, and evolutionary trends in leaf and flower morphology. Taxon 55(3), 617–642.

Kanehisa, M., and Goto, S. (2000). KEGG: Kyoto Encyclopedia of Genes and Genomes. Nucleic Acids Research 28(1), 27–30. doi:10.1093/nar/28.1.27.

Katschnig, D., Bliek, T., Rozema, J., and Schat, H. (2015). Constitutive high-level SOS1 expression and absence of HKT1; 1 expression in the salt-accumulating halophyte *Salicornia dolichostachya*. Plant Science 234, 144–154.

Khan, M.A., Ungar, I.A., and Showalter, A.M. (2000). The effect of salinity on the growth, water status, and ion content of a leaf succulent perennial halophyte, *Suaeda fruticosa* (L.) Forssk. Journal of Arid Environments 45(1), 73–84.

Kline, K.L., Msangi, S., Dale, V.H., Woods, J., Souza, G.M., Osseweijer, P., et al. (2017). Reconciling food security and bioenergy: priorities for action. Gcb Bioenergy 9(3), 557–576.

Li, X., Guo, C., Ahmad, S., Wang, Q., Yu, J., Liu, C., et al. (2019). Systematic Analysis of MYB Family Genes in Potato and Their Multiple Roles in Development and Stress Responses. Biomolecules 9(8), 317.

Livak, K.J., and Schmittgen, T.D. (2001). Analysis of relative gene expression data using Real-Time Quantitative PCR and the 2^-^ΔΔCT method. Methods 25(4), 402–408. doi: https://doi.org/10.1006/meth.2001.1262.

Loqué, D., Ludewig, U., Yuan, L., and von Wirén, N. (2005). Tonoplast intrinsic proteins *At*TIP2; 1 and *At*TIP2; 3 facilitate NH3 transport into the vacuole. Plant physiology 137(2), 671–680.

Madeira, F., Park, Y.m., Lee, J., Buso, N., Gur, T., Madhusoodanan, N., et al. (2019). The EMBL-EBI search and sequence analysis tools APIs in 2019. Nucleic Acids Research 47(W1), W636–W641. doi:10.1093/nar/gkz268.

Maurel, C., Boursiac, Y., Luu, D.-T., Santoni, V., Shahzad, Z., and Verdoucq, L. (2015). Aquaporins in plants. Physiological reviews 95(4), 1321–1358.

Mittler, R. (2006). Abiotic stress, the field environment and stress combination. Trends in plant science 11(1), 15–19.

Munns, R. (2002). Comparative physiology of salt and water stress. Plant, cell & environment 25(2), 239–250.

Munns, R., and Tester, M. (2008). Mechanisms of salinity tolerance. Annu. Rev. Plant Biol. 59, 651–681.

Negrão, S., Schmöckel, S.M., and Tester, M. (2016). Evaluating physiological responses of plants to salinity stress. Annals of Botany 119(1), 1–11. doi:10.1093/aob/mcw191.

Oh, D.-H., Hong, H., Lee, S.Y., Yun, D.-J., Bohnert, H.J., and Dassanayake, M. (2014). Genome structures and transcriptomes signify niche adaptation for the multiple-ion-tolerant extremophyte *Schrenkiella parvula*. Plant Physiology 164(4), 2123–2138.

Oh, M.-M., Carey, E.E., and Rajashekar, C. (2009). Environmental stresses induce health-promoting phytochemicals in lettuce. Plant Physiology and Biochemistry 47(7), 578–583.

Osakabe, Y., Osakabe, K., Shinozaki, K., and Tran, L.-S.P. (2014). Response of plants to water stress. Frontiers in plant science 5, 86–86. doi:10.3389/fpls.2014.00086.

Pandey, P., Irulappan, V., Bagavathiannan, M.V., and Senthil-Kumar, M. (2017). Impact of combined abiotic and biotic stresses on plant growth and avenues for crop improvement by exploiting physio-morphological traits. Frontiers in Plant Science 8(537). doi:10.3389/fpls.2017.00537.

Panta, S., Flowers, T., Lane, P., Doyle, R., Haros, G., and Shabala, S. (2014). Halophyte agriculture: success stories. Environmental and Experimental Botany 107, 71–83.

Parida, A.K., and Das, A.B. (2005). Salt tolerance and salinity effects on plants: a review. Ecotoxicology and environmental safety 60(3), 324–349.

Phukan, U.J., Jeena, G.S., and Shukla, R.K. (2016). WRKY transcription factors: molecular regulation and stress responses in plants. Frontiers in plant science 7, 760.

Platten, J.D., Cotsaftis, O., Berthomieu, P., Bohnert, H., Davenport, R.J., Fairbairn, D.J., et al. (2006). Nomenclature for HKT transporters, key determinants of plant salinity tolerance. Trends in plant science 11(8), 372–374.

Pucker, B., and Schilbert, H.M. (2019). “Genomics and Transcriptomics Advance in Plant Sciences,” in Molecular Approaches in Plant Biology and Environmental Challenges, eds. S.P. Singh, S.K. Upadhyay, A. Pandey & S. Kumar. (Singapore: Springer Singapore), 419–448.

Qadir, M., Quillérou, E., Nangia, V., Murtaza, G., Singh, M., Thomas, R.J., et al. (Year). “Economics of salt-induced land degradation and restoration”, in: Natural resources forum: Wiley Online Library), 282–295.

Ramegowda, V., and Senthil-Kumar, M. (2015). The interactive effects of simultaneous biotic and abiotic stresses on plants: Mechanistic understanding from drought and pathogen combination. Journal of Plant Physiology 176, 47–54. doi: https://doi.org/10.1016/j.jplph.2014.11.008.

Ray, D.K., Mueller, N.D., West, P.C., and Foley, J.A. (2013). Yield trends are insufficient to double global crop production by 2050. PloS one 8(6), e66428.

Riechmann, J.L., Heard, J., Martin, G., Reuber, L., Jiang, C.-Z., Keddie, J., et al. (2000). *Arabidopsis* transcription factors: genome-wide comparative analysis among eukaryotes. Science 290(5499), 2105–2110.

Rivero, R.M., Mestre, T.C., Mittler, R., Rubio, F., Garcia-Sanchez, F., and Martinez, V. (2014). The combined effect of salinity and heat reveals a specific physiological, biochemical and molecular response in tomato plants. Plant, Cell & Environment 37(5), 1059–1073. doi:10.1111/pce.12199.

Roy, S.J., Negrão, S., and Tester, M. (2014). Salt resistant crop plants. Current opinion in Biotechnology 26, 115–124.

Rozema, J., and Schat, H. (2013). Salt tolerance of halophytes, research questions reviewed in the perspective of saline agriculture. Environmental and Experimental Botany 92, 83–95.

Rushton, P.J., Somssich, I.E., Ringler, P., and Shen, Q.J. (2010). WRKY transcription factors. Trends in plant science 15(5), 247–258.

Salazar, O. (2017). Identification of proteins involved in salinity tolerance in Salicornia bigelovii. PhD, King Abdullah University of Science and Technology.

Shao, Q., Zhao, C., Han, N., and Wang, B.-S. (2008). Cloning and expression pattern of *Ss*HKT1 encoding a putative cation transporter from halophyte *Suaeda salsa*. DNA Sequence 19(2), 106–114.

Shi, H., Ishitani, M., Kim, C., and Zhu, J.-K. (2000). The *Arabidopsis thaliana* salt tolerance gene SOS1 encodes a putative Na^+^/H^+^ antiporter. Proceedings of the national academy of sciences 97(12), 6896–6901.

Shukla, P.S., Agarwal, P., Gupta, K., and Agarwal, P.K. (2015). Molecular characterization of an MYB transcription factor from a succulent halophyte involved in stress tolerance. AoB plants 7.

Silva, P., and Gerós, H. (2009). Regulation by salt of vacuolar H^+^-ATPase and H^+^-pyrophosphatase activities and Na^+^/H^+^ exchange. Plant Signaling & Behavior 4(8), 718–726. doi:10.4161/psb.4.8.9236.

Singh, K.B., Foley, R.C., and Oñate-Sánchez, L. (2002). Transcription factors in plant defense and stress responses. Current opinion in plant biology 5(5), 430–436.

Singh, L.P., Gill, S.S., and Tuteja, N. (2011). Unraveling the role of fungal symbionts in plant abiotic stress tolerance. Plant signaling & behavior 6(2), 175–191.

Song, J., and Wang, B. (2015). Using euhalophytes to understand salt tolerance and to develop saline agriculture: *Suaeda salsa* as a promising model. Annals of Botany 115(3), 541–553.

Statistics Center (2018). “Statistical Yearbook of Abu Dhabi 2017”. Statistics Centre of Abu Dhabi).

Su, H., Balderas, E., Vera-Estrella, R., Golldack, D., Quigley, F., Zhao, C., et al. (2003). Expression of the cation transporter McHKT1 in a halophyte. Plant Molecular Biology 52(5), 967–980.

Tang, Y., Bao, X., Zhi, Y., Wu, Q., Yin, X., Zeng, L., et al. (2019). Overexpression of a MYB family gene, OsMYB6, increases drought and salinity stress tolerance in transgenic rice. Frontiers in plant science 10, 168.

The Gene Ontology Consortium (2018). The Gene Ontology Resource: 20 years and still GOing strong. Nucleic Acids Research 47(D1), D330–D338. doi:10.1093/nar/gky1055.

UN (2019). “World population prospects 2019”. United Nations).

Vaishnav, A., Shukla, A.K., Sharma, A., Kumar, R., and Choudhary, D.K. (2019). Endophytic bacteria in plant salt stress tolerance: Current and future prospects. Journal of Plant Growth Regulation 38(2), 650–668. doi:10.1007/s00344-018-9880-1.

Van Aken, O., Zhang, B., Law, S., Narsai, R., and Whelan, J. (2013). AtWRKY40 and AtWRKY63 modulate the expression of stress-responsive nuclear genes encoding mitochondrial and chloroplast proteins. Plant physiology 162(1), 254–271.

Wang, G.-L., Tian, C., Jiang, Q., Xu, Z.-S., Wang, F., and Xiong, A.-S. (2016). Comparison of nine reference genes for real-time quantitative PCR in roots and leaves during five developmental stages in carrot (*Daucus carota* L.). The Journal of Horticultural Science and Biotechnology 91(3), 264–270. doi:10.1080/14620316.2016.1148372.

Wang, S.-P., Krits, I., Bai, S., and Lee, B.S. (2002). Regulation of enhanced vacuolar H^+^-ATPase expression in macrophages. Journal of Biological Chemistry 277(11), 8827–8834.

Wang, S., DUan, L.W., Jl, Z., and Sm, W. (2013). “Molecular cloning and sequence analysis of plasma membrane Na^+^/H^+^ antiporter gene (SsSOS1) from halophyte Suaeda salsa”.).

Warshay, B., Brown, J.J., and Sgouridis, S. (2017). Life cycle assessment of integrated seawater agriculture in the Arabian (Persian) Gulf as a potential food and aviation biofuel resource. The International Journal of Life Cycle Assessment 22(7), 1017–1032.

Wu, Y., Hu, Y., and Xu, G. (2009). Interactive effects of potassium and sodium on root growth and expression of K/Na transporter genes in rice. Plant Growth Regulation 57(3), 271.

Yuan, F., Guo, J., Shabala, S., and Wang, B. (2019). Reproductive physiology of halophytes: current standing. Frontiers in plant science 9, 1954.

Zerai, D.B., Glenn, E.P., Chatervedi, R., Lu, Z., Mamood, A.N., Nelson, S.G., et al. (2010). Potential for the improvement of *Salicornia bigelovii* through selective breeding. Ecological Engineering 36(5), 730–739.

Zhang, G.-H., Su, Q., An, L.-J., and Wu, S. (2008). Characterization and expression of a vacuolar Na^+^/H^+^ antiporter gene from the monocot halophyte *Aeluropus littoralis*. Plant Physiology and Biochemistry 46(2), 117–126.

Zhao, L., Yang, Z., Guo, Q., Mao, S., Li, S., Sun, F., et al. (2017). Transcriptomic profiling and physiological responses of halophyte *Kochia sieversiana* provide insights into salt tolerance. Frontiers in Plant Science 8(1985). doi: 10.3389/fpls.2017.01985.

Zhu, J.-K. (2002). Salt and drought stress signal transduction in plants. Annual Review of Plant Biology 53(1), 247–273. doi: 10.1146/annurev.arplant.53.091401.143329.

Zörb, C., Geilfus, C.M., and Dietz, K.J. (2019). Salinity and crop yield. Plant biology 21, 31–38.

