## Supplementary Data for "Beyond the greenhouse: coupling environmental and salt stress response reveals unexpected global transcriptional regulatory networks in *Salicornia bigelovii*"

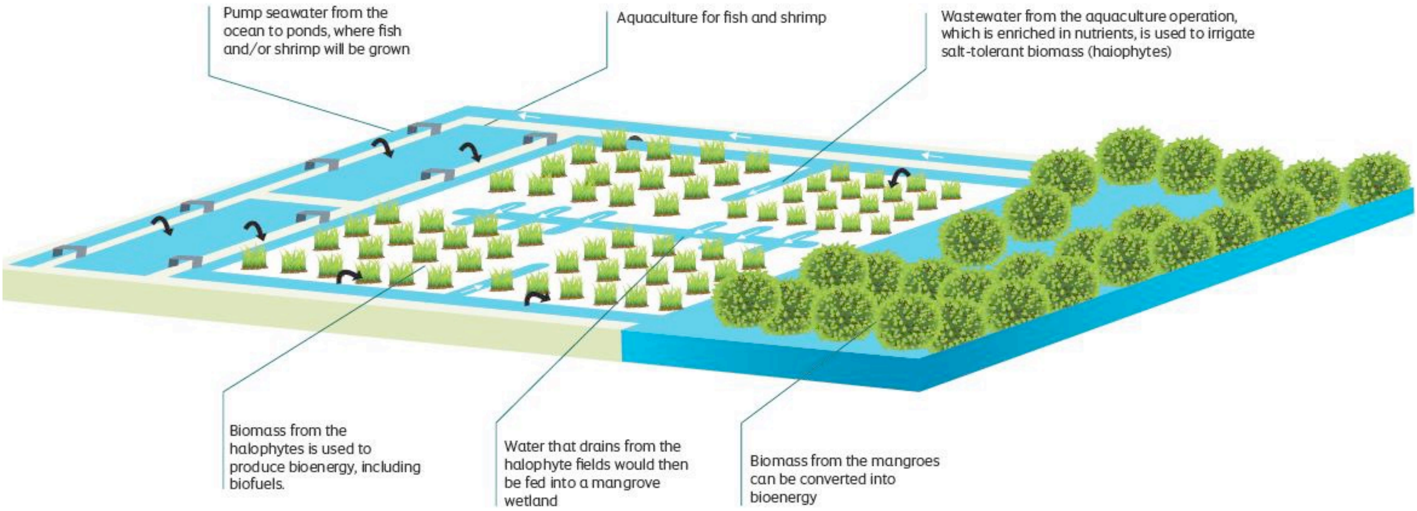


**Figure S1**. Schematic diagram of the SEAS pilot facility at Masdar Campus, Khalifa University. The pilot facility combines aquaculture, halo-agriculture, and mangrove silviculture for the production of sustainable biofuels for aviation and other high value byproducts such as seafood and animal fodder.

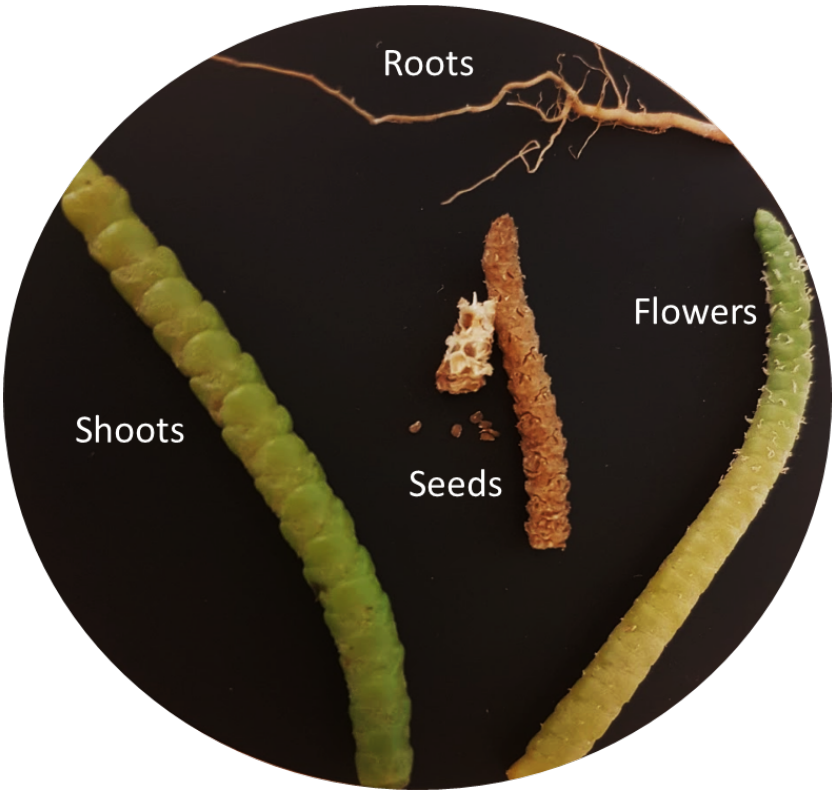


**Figure S2.** Image showing the different tissues used in the *S. bigelovii* transcriptome analysis. The *S. bigelovii* plant roots extend 20-35 cm into the soil, shoot height ranges from 9-60 cm, flowers are 0.6-0.7 mm, and seeds 1-1.5 mm in length.

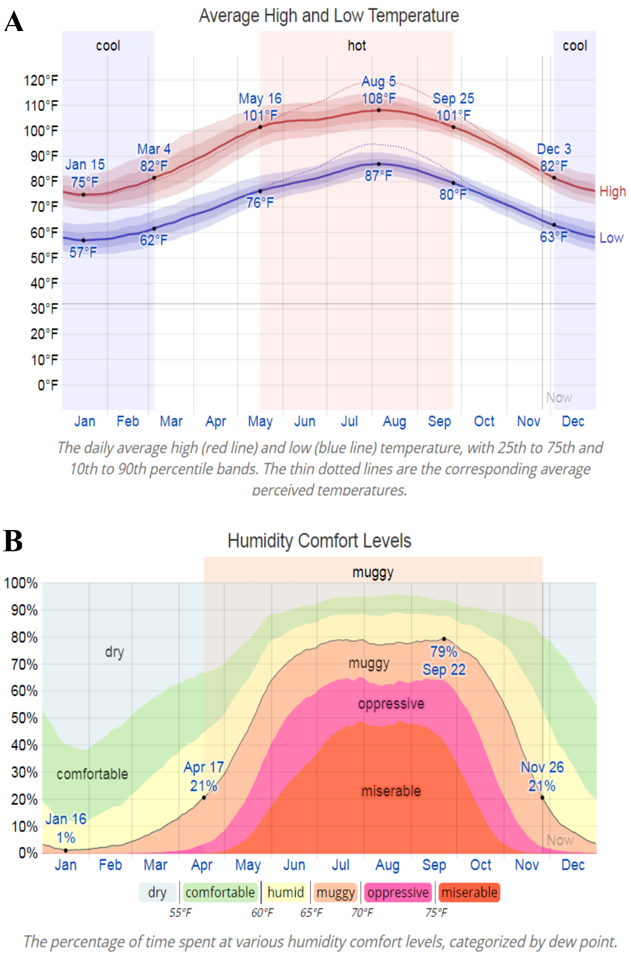


**Figure S3**. Meteorological data at the SEAS pilot facility as reported by the Abu Dhabi International Airport (24°25'59"N 54°39'04"E). (**A**) Graph represents the average monthly high (red line) and low (blue line) temperatures. (**B**) Image represents the monthly relative humidity levels with average time spent at various comfort levels.

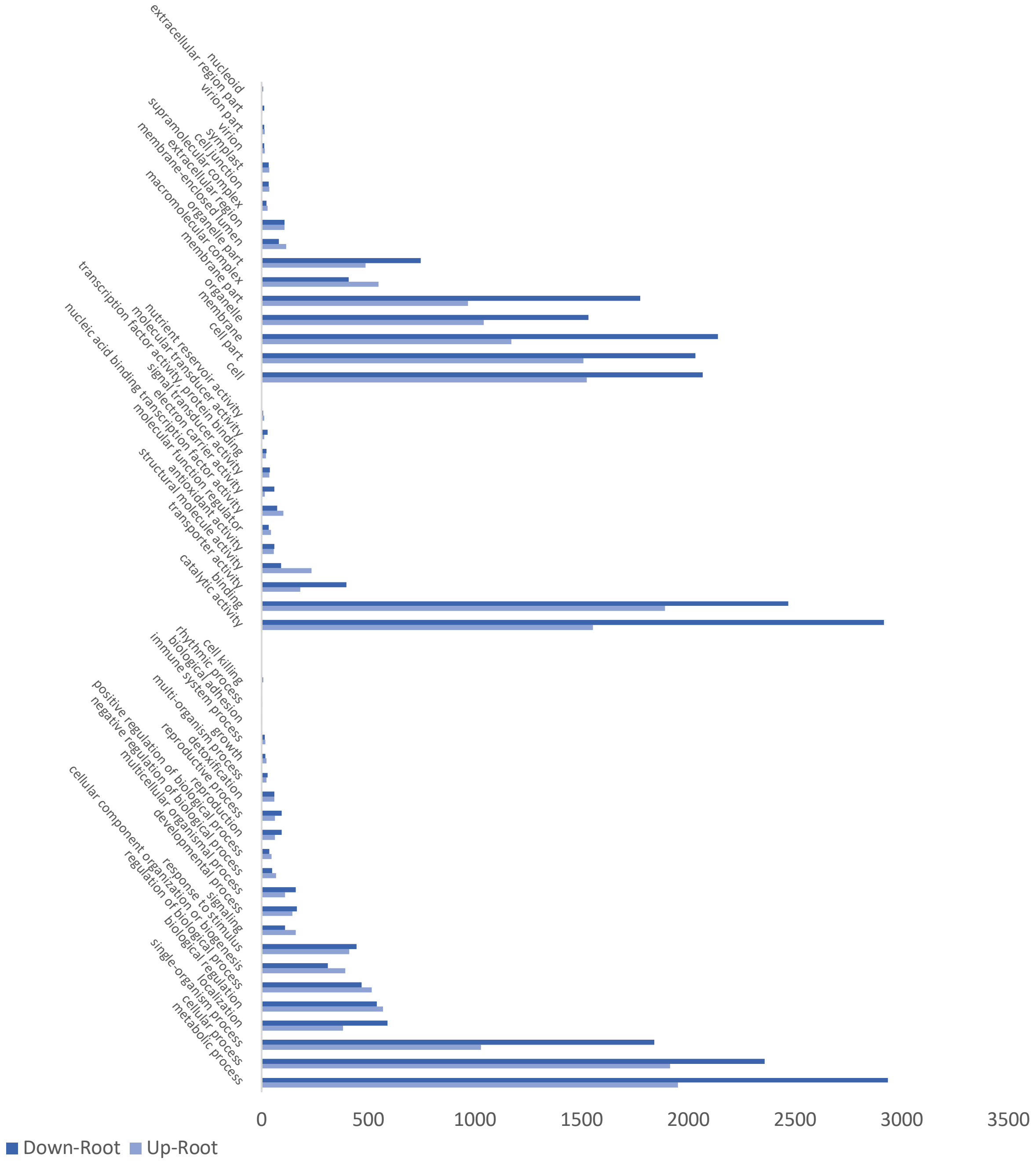


**Figure S4.** Gene Ontology classification of DETs root vs shoot. GO categories shown in the Y axis are grouped into three main ontologies, (A) cellular components (CC), (B) molecular functions (MF), and (C) biological processes (BP). The X axis represents the total number of transcripts down regulated (dark blue) and upregulated (light blue) in root tissue.

**Table S1.** Salinity and ion concentration of the effluent water in the Field 1 of SEAS pilot facility measured between March and August 2017.

| **Ions** | **March ($g\ L^{-1}$)** | **April ($g\ L^{-1}$)** | **May ($g\ L^{-1}$)** | **June ($g\ L^{-1}$)** | **July ($g\ L^{-1}$)** | **August ($g\ L^{-1}$)** | **Average ($g\ L^{-1}$)** |
|---|---|---|---|---|---|---|---|
| Na | 11.77 | 14.57 | 14.57 | 16.24 | 16.63 | 16.97 | 15.12 ±1.9 |
| Cl | 18.79 | 20.47 | 19.47 | 21.98 | 21.77 | 22.00 | 20.74 ±1.3 |
| Mg | 1.16 | 1.24 | 1.24 | 1.35 | 1.35 | 1.42 | 1.29 ±0.09 |
| Ca | 0.65 | 0.70 | 0.70 | 0.73 | 0.73 | 0.83 | 0.72 ±0.06 |
| K | 0.53 | 0.52 | 0.52 | 0.54 | 0.54 | 0.66 | 0.55 ±0.05 |
| Br | 0.06 | 0.06 | 0.06 | 0.07 | 0.07 | 0.09 | 0.07 ±0.01 |
| SO4 | 0.31 | 0.32 | 0.32 | 0.38 | 0.38 | 0.39 | 0.35 ±0.03 |
| NO2 | 0.00 | 0.00 | 0.00 | 0.01 | 0.01 | 0.01 | 0.01 ±0.02 |
| Water Salinity | 34.57 ±0.76 | 38.59 ±0.15 | 36.21 ±0.79 | 41.45 ±0.83 | 41.74 ±0.75 | 42.56 ±1.1 | |

**Table S2.** Primer sense and anti-sense primer sequences used in the qRT-PCR assay for validation of differential expression of specific genes.

| Transcript ID | Sequence(5'to3') | Reverse | Description |
|---|---|---|---|
| TRINITY_DN119119 | CGAACCTTCGAAGATGGAGA | CTTATTTCGTGTGCCGGTTT | *Beta vulgaris* subsp. *vulgaris* probable cation transporter HKT6 |
| TRINITY_DN220726 | CGAACCTTCGAAGATGGAGA | CTTATTTCGTGTGCCGGTTT | *Salicornia brachiata* salt overly sensitive 1 mRNA, complete CDS |
| TRINITY_D N207378 | TGTACCAATGGCTCCAAACA | GGGGTTGTGATTCTGCTGAT | *Salicornia bigelovii* Na+/H+ antiporter (NHX1) mRNA, complete CDS |
| TRINITY_DN190896 | CCTCTCTTCCTTGGCTTCCT | ACTTTGGTGAGGTTGGTGCT | *Chenopodium quinoa* abscisic stress-ripening protein 3-like, mRNA |
| TRINITY_DN98123 | CCCCTCCACAGTGTTCCTTA | ACAAACCAAGCCACACAACA | *Beta vulgaris* subsp. *vulgaris* probable aquaporin NIP 5, mRNA |
| TRINITY_DN231325 | AGCACCAGCTTCTGGACCTA | TTCAAAAGCCATGTGCTCAG | *Kalidium foliatum* betaine aldehyde dehydrogenase protein mRNA, complete CDS |
| TRINITY_DN218725 | CACCCCAATAACATCGGAAC | TGACGACAACGACGAAGAAG | Proline-rich protein PRCC (LOC104883603), transcript variant X2, mRNA |
| TRINITY_DN223373 | TCGCAGCAAGAGAGAGTTCA | CGTATCCAGCTTGTTGAGCA | *Beta vulgaris* subsp. *vulgaris* proline-rich protein P5CS |
| TRINITY_DN206459 | TGAATCATCCTGTGCTGCTC | CTGGCTCAATCTCGTTGACA | *Salicornia europaea* vacuolar H+-pyrophosphatase mRNA, complete CDS |
